# Sugar assimilation underlying dietary evolution of Neotropical bats

**DOI:** 10.1101/2023.07.02.547432

**Authors:** Jasmin Camacho, Andrea Bernal-Rivera, Valentina Peña, Pedro Morales-Sosa, Sofia Robb, Jonathon Russell, Kexi Yi, Yongfu Wang, Dai Tsuchiya, Oscar E. Murillo-García, Nicolas Rohner

## Abstract

Specializations in animal diets drive selective demands on morphology, anatomy, and physiology. Studying adaptations linked to diet evolution benefits from examining Neotropical bats, a remarkable group with high taxonomic and trophic diversity. In this study, we performed glucose tolerance tests on wild-caught bats, which revealed distinct responses to three sugars present in different foods: trehalose (insects), sucrose, and glucose (fruits and nectar). Insect-eating bats metabolism responded most strongly to trehalose, while bats with nectar and fruit-based diets exhibited a heightened response to glucose and sucrose, reaching blood glucose levels over 600 and 750 mg/dL. To search for signatures of positive selection in sugar assimilation genes we performed genome analysis of 22 focal bat species and 2 outgroup species. We identified selection in the ancestral vespertilionid branch (insect-eaters) for the digestive enzyme trehalase, while sucrase-isomaltase exhibited selection in branches leading to omnivorous and nectar diets. Unexpectedly, the insect-eating lineage *Myotis* exhibited sucrase-isomaltase selection, potentially explaining their heightened sucrose assimilation. Furthermore, the genes encoding for glucose transporters, *Slc2a3* and *Slc2a2,* showed selection in nectar and blood feeding bats, with analyses of predicted protein structures supporting modified activity. By examining cellular features of the small intestine, we discovered that dietary sugar proportion strongly impacted numerous digestive traits, providing valuable insight into the physiological implications of the identified molecular adaptations. To elucidate this further, we used HCR RNA-FISH to perform single molecule *ex vivo* gene expression analysis of enterocyte response to a glucose meal in three focal species. We observed unusually high activity in the glucose transporter *Slc2a2* during the fasted state of nectar bats that did not change upon feeding. Comparatively, nectar bats exhibited an enhanced capacity for intestinal absorption of dietary sugar primarily through *Slc2a2*, while fruit bats relied on increasing levels of *Slc5a1*. Overall, this study highlights the intricate interplay between molecular, morphological, and physiological aspects of diet evolution and provides new insights into our understanding of dietary diversification and sugar assimilation mechanisms in mammals.

**Graphical Abstract:** 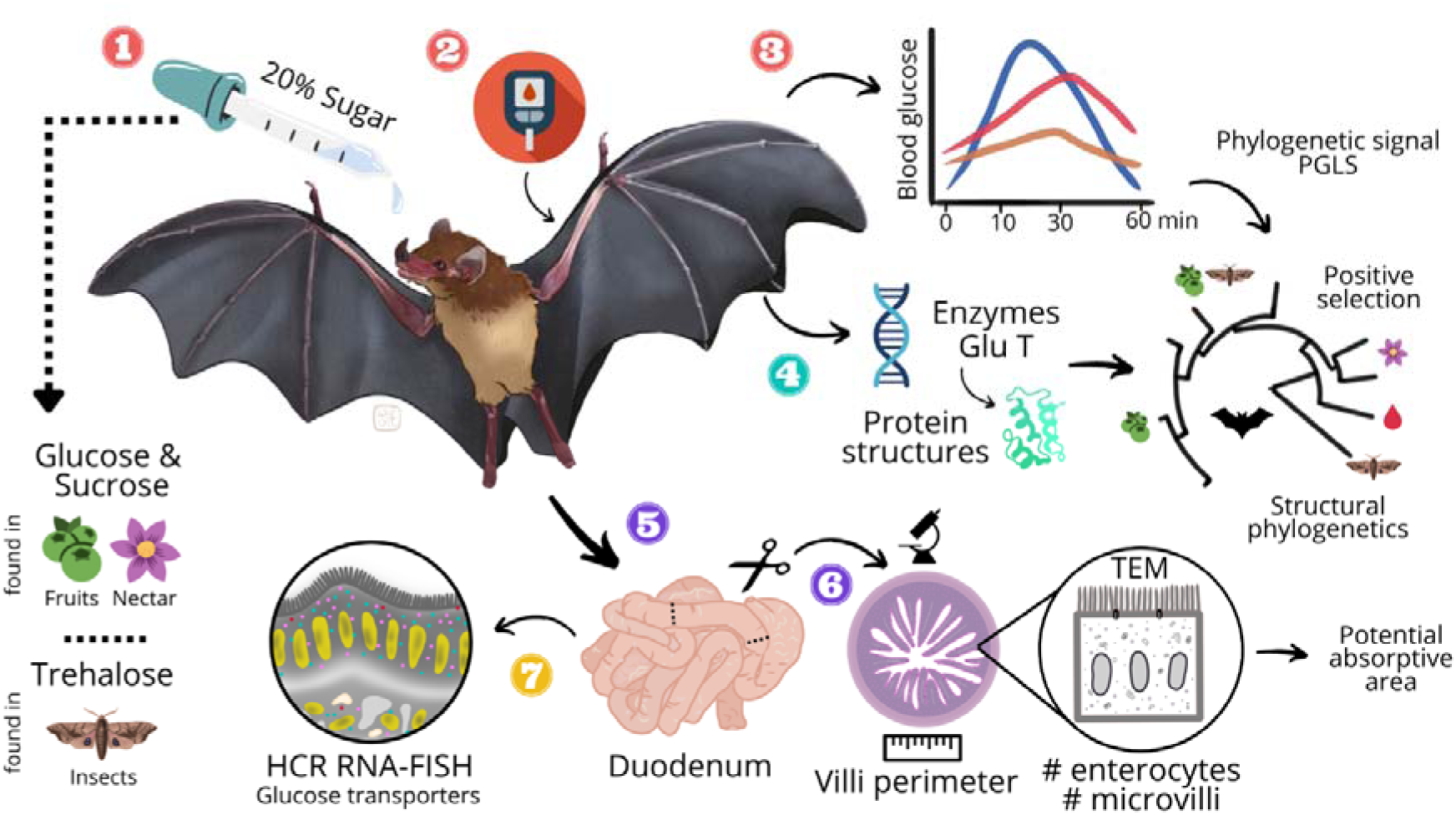

**Highlights:**

- Sugar assimilation differences emphasize metabolic adaptations to diet
- Glucose tolerance tests provide a quick and practical assessment of dietary ecology
- Bat genomes exhibit positive selection on digestive enzymes and glucose transporters
- Structural comparisons of proteins suggest altered activity of glucose transporters
- Glucose absorption differences can be explained by gut anatomy
- Intestinal villus diversity and novel microanatomy in bats
- Extreme blood glucose (above 600 and 750 mg/dL) coincides with constitutive expression of apical *Slc2a2*
- The regulation of apical *Slc2a2* highlights differences in blood glucose levels

## Introduction

Organisms actively adapt to their environments to improve their chances of survival and reproductive success. Throughout animal evolution, diet is a major environmental input that influences adaptations in nutrient acquisition, such as feeding morphology and behavior (Pauli et al. 2016; Tokita et al. 2017; Camacho et al. 2019; Morris et al. 2019; Lyons et al. 2020; Dobreva et al. 2022; Yohe et al. 2022; Sadier et al. 2023). Changes in metabolism underlie these diet-related evolutionary variations, affecting the ability of cells and tissues to sense and respond to altered nutrient availability. This ability is crucial for maintaining energy homeostasis and overall health (Deas et al. 2019; Koyama et al. 2020; Karasov & Caviedes-Vidal, 2021; Riddle et al. 2021; Xiong 2022).

Glucose homeostais is the central mechanism that governs environmental adaptation to nutritional sources, with the intestinal tract villi being crucial in this process since many nutrients are digested and absorbed here. Glucose homeostasis is a tightly regulated biochemical pathway that maintains circulating glucose levels within a narrow physiological range for cellular function and organismal energy. Blood glucose levels fluctuate throughout the day, are easily measured, and relate to lifestyle and diet (Fujisaka et al. 2018). This pathway is expected to have a critical role in metabolic adaptations (Karasov & Douglas, 2013). However, wildlife studies investigating these adaptations are scarce and include few taxa (Costa & Ortiz, 1982; Tomasek et al. 2018; Sparkman et al. 2018; Bennett et. al. 2017; Saxton et al. 2022). Existing research is primarily focused on accessible lab populations of wild animals (Oriel et al. 2008; Kelm et al. 2011; Riddle et al. 2018; Reznick et al. 2021; Guo et al. 2022) or restricted within the context of metabolic disorders (Chouchani & Kajimura 2019; Bundgaard et al. 2019).

Natural systems exhibiting a wide diversity of diets, such as bats, provide a unique opportunity to investigate diet-related evolutionary changes. Bats have diversified from an insect-heavy ancestral diet to nutritional sources including fruit, nectar, meat, and fish, among others (Neuweiler, 2000; Arbour 2019). The dietary shifts among bats reflect the evolutionary changes across major clades of mammals (Price 2012; Camacho et al. 2020), such as herbivores, omnivores, and carnivores (i.e. Orders Artiodactyla, Primates, Carnivora). Therefore, investigating their adaptive radiation is expected to provide new perspectives on the diversification of metabolic traits and emphasize the molecular basis of these adaptations.

In this study, we explore the metabolic adaptations involved in dietary changes. We observe adaptation to the glucose homeostasis pathway across 29 bat species with different diets, using three dietary sugars-trehalose (found in insects’ hemolymph), sucrose, and glucose (both found in fruits and nectar). We demonstrate that the metabolic phenotype is mediated by four different adaptations to digestive morphology, including intestinal length, exposed villi, and enterocyte and microvilli number. We have discovered genetic traits linked to maximal extraction of glucose energy from nutritional resources associated with diet. These include positive selection on genes encoding the digestive enzymes Trehalase (*Treh*) and Sucrase-Isomaltase (*SI*), as well as glucose transporters (*Slc2a1-5*). We investigated the effects of these amino acid substitutions on protein function through structural comparisons. Additionally, we observed a shift in the expression of genes encoding transporters SLC2A2 (*Slc2a2*), SCL2A5 (*Slc2a5*), and SLC5A1 (*Slc5a1*) within enterocytes along the brush border of the small intestine. The emerging picture illustrates the sophisticated adaptability of absorptive villi, highlighting the evolution of traits and mechanisms that enable species to utilize a broader range of nutrient resources compared to those found ancestrally. Overall, this study advances our understanding of how metabolic evolution contributes to species diversification.

## Results and Discussion

### *In vivo* physiology

Among mammals, bats offer a unique opportunity to investigate metabolic adaptations due to their high energetic demands and a wide variety of diets. They have specialized from an ancestor with an insect-heavy diet (Simmons et al., 2008) to fruit, nectar, meat, blood, and other food sources (Neuweiler, 2000). To assess metabolic adaptations related to dietary diversification, we focused on the glucose homeostasis pathway, which directly regulates glucose energy during and between food intake and is vital for long-term health and survival. We performed *in vivo* oral glucose tolerance tests on 199 wild-caught bats from 29 species (Supplemental Table 1) to assess their ability to process three dietary sugars (Wyatt & Kalf, 1957; Baker et al., 1998): trehalose (formed by two glucose molecules), sucrose (formed by glucose and fructose), and glucose, expecting higher assimilation of the absorbable monosaccharide compared to the more complex sugars which undergo hydrolysis before being absorbed. The bats were fasted for 10-12 hours, and blood glucose levels were measured before and after consuming a single sugar meal (5.4g/kg, Kelm et al., 2011). Measurements were taken at 10, 30, and 60 minutes post-ingestion, revealing distinct patterns in the rise and fall of blood glucose levels between diets (Fig. 1A). To better quantify physiological responses, we classified them into four assimilation patterns: *“Fast”*, *“Medium”*, *“Slow”*, and *“Limited”* (Fig. 1B), based on the speed and extent of assimilation (see Methods 1.2.1).

**Fig. 1.**
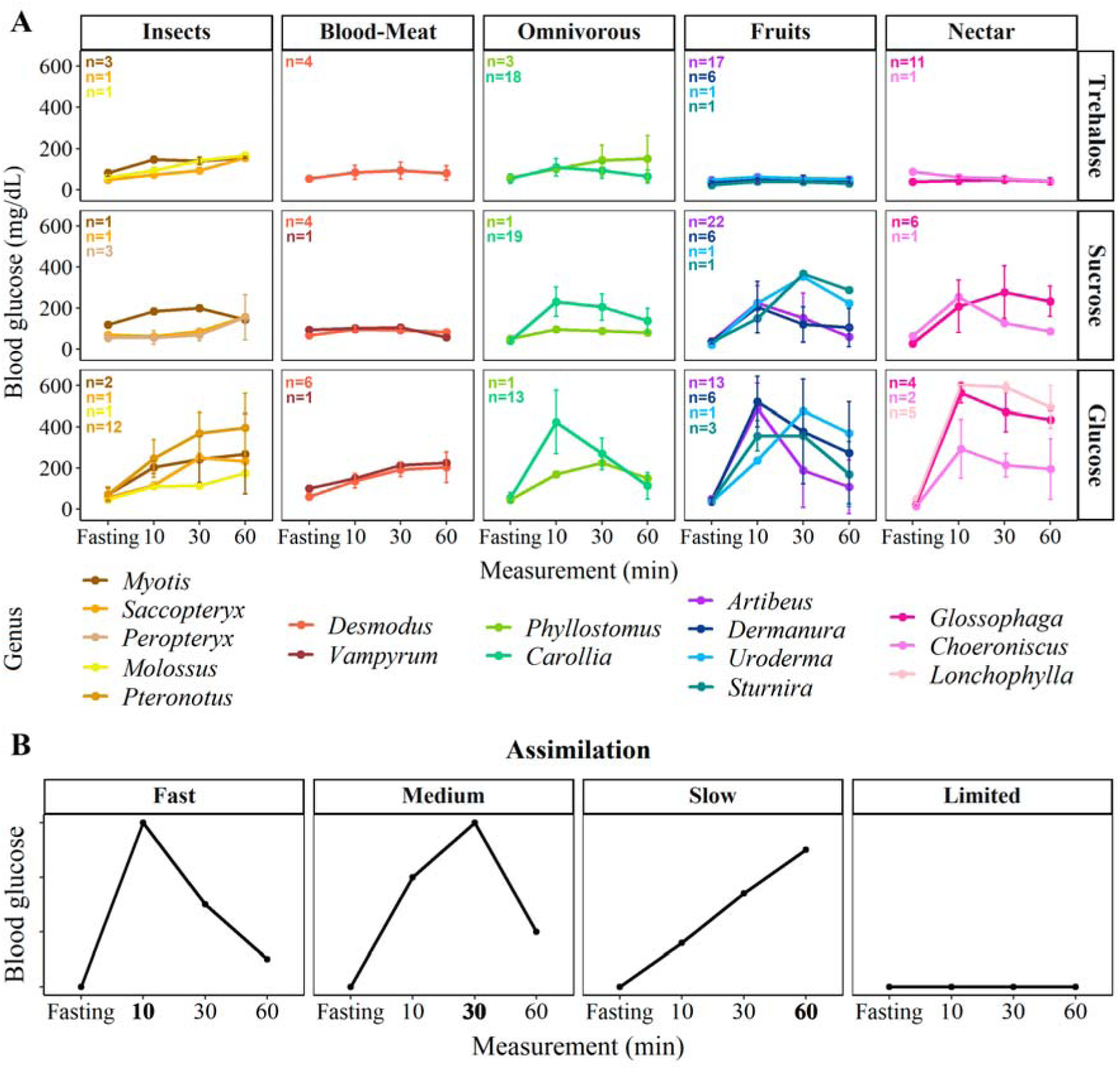
A. Average assimilation curves for trehalose, sucrose, and glucose among Neotropical bats with different food preferences: insects, blood or meat, mixed (omnivorous), fruits, and nectar. The sample size varied among genera from 1 to 52 individuals (Supplemental Table 1). B. General curves to describe the temporal pattern of sugar assimilation.

In insectivorous and omnivorous bats, we observed <u>trehalose</u> assimilation with blood glucose levels reaching up to 160 mg/dL with a *“Slow”* tolerance curve indicating glucose absorption continuing until 60 minutes. In contrast, bats with nectar and fruit diets exhibited a *“Limited”* ability to assimilate trehalose, showing blood glucose levels remaining within a narrow range and only rising to 60 mg/dL. These findings align with existing evidence that the trehalase gene (*Treh*) is non-functional in non-insectivorous mammals, including bats, at the genome and protein level, and functional in omnivores such as *Carollia* and *Phyllostomus* (Schondube et al., 2001; Jiao et al., 2019). Interestingly, individuals of the vampire bat showed a slight rise and fall in blood glucose levels following trehalose consumption, reaching levels of 90 mg/dL. We propose that the minor elevation in blood glucose levels in non-insectivorous bats could be attributed to the activity of the gut microbiome (Zepeda Mendoza et al., 2018; Ingala et al., 2021), as trace amounts of trehalase was found in their gut (Schondube et al., 2001).

When tested for <u>sucrose</u>, insect-feeding bats showed a gradual increase in blood glucose levels 30 minutes after sucrose ingestion, except for *Myotis*, which surprisingly showed a rapid rise to 200 mg/dL within 10 minutes, followed by a decline to 140 mg/dL at 60 minutes, indicating a “*Medium*” sucrose assimilation. This suggests that *Myotis* might be capable of sucrose digestion more efficiently than trehalose despite its classification as an insect feeder. As expected, the vampire bats showed *“Limited”* sucrose assimilation, with blood glucose levels remaining below 96 mg/dL throughout the experiment. Among omnivores, *Carollia* levels peaked at 10 min (200 mg/dL) and gradually declined, whereas *Phyllostomus* had a lower peak at 10 min (100 mg/dL), and slowly decreased. These patterns reflect the varying degrees of omnivory, where *Carollia* mainly feed on piper fruits and some insects, while *Phyllostomus* consume insects, fruits, small vertebrates, flowers, nectar and pollen (Santos et al., 2003; York & Billings, 2009). Fruit and nectar bats exhibited two major responses to sucrose: 1. “*Fast”* assimilation with a rapid rise in blood glucose (200 mg/dL) within 10 minutes and subsequent rapid decrease to basal levels after 60 min for *Artibeus, Dermanura,* and *Choeroniscus*; 2. “*Medium”* assimilation with a slow increase, reaching a maximum after 30 minutes (300 mg/dL) for *Sturnira, Uroderma,* and *Glossophaga.* These findings highlight significant variations in sucrose assimilation among bats with different diets, and within species of the same dietary category, indicating the diversification of mechanisms for disaccharide assimilation and probably seasonality impact that is worth analyzing further, especially for omnivorous bats.

When tested for <u>glucose</u> ingestion, we observed higher blood glucose levels across all bat species compared to trehalose and sucrose feedings. This is because monosaccharides can be directly absorbed through the intestine while disaccharides (sucrose and trehalose) require previous hydrolysis (Price et al., 2015). Bats with insect, meat, and blood diets showed a gradual increase in blood glucose levels and stayed below 300 mg/dL, except in *Pteronotus* (> 350 mg/dL), but still indicating a *“Slow”* assimilation pattern. Among omnivorous bats, *Carollia* reached a peak of 423 mg/dL at 10 minutes and rapidly decreased levels until 60 minutes, while *Phyllostomus* reached a peak of 225 mg/dL at 30 minutes and then declined by 60 minutes. Bats with fruit diets exhibited the most extreme rise and fall of blood glucose levels, with levels above 600 mg/dL within 10 minutes in *Artibeus* and *Dermanura*, and levels dropping to 100 mg/dL in some individuals after 60 minutes. This *“Fast”* assimilation curve is likely due to an enhanced insulin sensitivity in these species (Protzek et al., 2010). Other fruit bats showed high glucose levels within 30 minutes, with *Uroderma* reaching 476 mg/dL and *Sturnira* reaching 357 mg/dL, followed by a subsequent decrease. In nectarivores, we observed *“Fast”* assimilation curves, with *Glossophaga* and *Lonchophylla* reaching levels above the detection limit of 600 mg/dL (GlucoQuick G30a) at 10 minutes and *Choeroniscus* reaching 300 mg/dL at the 10-minute time point, which is probably related to differences in plant/nectar preferences and food availability for each species (Ayala-Berdon et al., 2013; Ortega-García et al., 2022). In contrast, captive *Glossophaga soricina* individuals reached a peak of 360 mg/dL after 30 minutes when fed with a similar single dose (Kelm et al., 2011) and did not exceed 470 mg/dL even when fed a higher glucose dose (9 g/kg). Nectar bats from Glossophaginae and Lonchophyllinae subfamilies exhibited a slower decline in blood glucose levels compared to fruit bats, consistent with recent studies that found no plasma insulin response to feeding in nectar-bats at rest and proposed an exercise-mediated regulation of glucose homeostasis (Welch et al., 2008; Welch & Chen, 2014; Kelm et al., 2011; Peng et al., 2017; Castro et al., 2021). These findings highlight how sugar assimilation capability in Neotropical bats reflects their natural food preferences, raising questions about how bats with different diets can absorb and compensate for variations in circulating blood glucose.

To evaluate the impact of evolutionary history on sugar assimilation, we employed Pagel’s λ (Pagel, 1999) to test the phylogenetic signal for each assimilation pattern with the area under the glucose tolerance test curve as a proxy. We obtained λ values of 0.99 for sucrose assimilation (p<0.05) (Supplemental information Table 2). However, glucose and trehalose did not show a phylogenetic signal. These results suggest that phylogenetic relationships are probably playing an important role in determining sucrose assimilation, but not in the monosaccharide or trehalose assimilation, which is likely why we see most variation among species in response to glucose and the higher response to trehalose in insectivorous and omnivorous bats even when they are not closely related. This observation is consistent with a prior study that examined traits related to digestion and found that the gut surface area, body mass, and digestion is not constrained by phylogeny, demonstrating its potential for adaptation (Saldaña_Vázquez et al., 2015). Additionally, we fitted a Bayesian multilevel phylogenetic model to compare adaptations in glucose assimilation while controlling for phylogenetic relatedness. The results showed differences in the assimilation pattern of glucose, sucrose, and trehalose between species with high and low sugar content in their diets (sucrose and glucose being higher in fruits and nectar, and trehalose being higher in insects). However, the omnivorous bats, with the three sugars being in relatively high proportions in their diets, showed similar glucose and sucrose assimilation patterns to bats with rich glucose and sucrose diets (fruit and nectar bats), and similar trehalose assimilation patterns to bats with rich trehalose diets (insectivorous) (Supplemental information Figure 5). This indicates a strong association between the assimilation patterns and dietary preferences.

### Molecular adaptations related to sugar assimilation

To identify molecular adaptations underlying the association between sugar assimilation and different diets, we focused on genes involved in glucose transport and absorption, crucial for regulating blood glucose levels and facilitating glucose uptake into cells (Figure 2). We tested for positive selection in 22 bats represented in our *in vivo* data, along with two outgroup species (Supplemental Table 3). We focused We analyzed the genes encoding the digestive enzyme trehalase (*Treh*), responsible for breaking down trehalose into two glucose molecules, and the enzyme sucrase-isomaltase (*SI*), which breaks down sucrose into one molecule of glucose and one of fructose in the small intestine (Figure 2A). We also examined genes encoding transporters involved in the absorption of these sugars from the intestinal lumen and into the blood, including the sodium/glucose cotransporter (*Slc5a1*), the solute carrier family 2 member 2 (*Slc2a2*), and the solute carrier family 2 member 5 (*Slc2a5*). Interestingly, while SLC2A5/GLUT5 functions primarily as a transporter for fructose, it possesses the capability to transport glucose (Burant et al., 1992; Mizuno et al., 2021). Furthermore, glucose transporters are known to play crucial roles in sequestering circulating blood glucose into cells to meet tissue-specific energy demands (Navale & Paranjape, 2016; Figure 2B). These genes include *Slc2a1*, with enriched expression in red blood cells and astrocytes (Veys et al. 2020), *Slc2a2*, enriched in the intestine, liver, hypothalamus, and pancreatic beta cells (Thorens 2015), *Slc2a3*, predominantly found in neurons (Gerhart et al. 1992), *Slc2a4*, expressed in insulin-sensitive tissues such as cardiac muscle, skeletal muscle, and fat (Richter and Hargreaves 2013), and *Slc2a5*, enriched in microglia, liver, kidney, and testis (Burant et al. 1992).

**Fig. 2.**
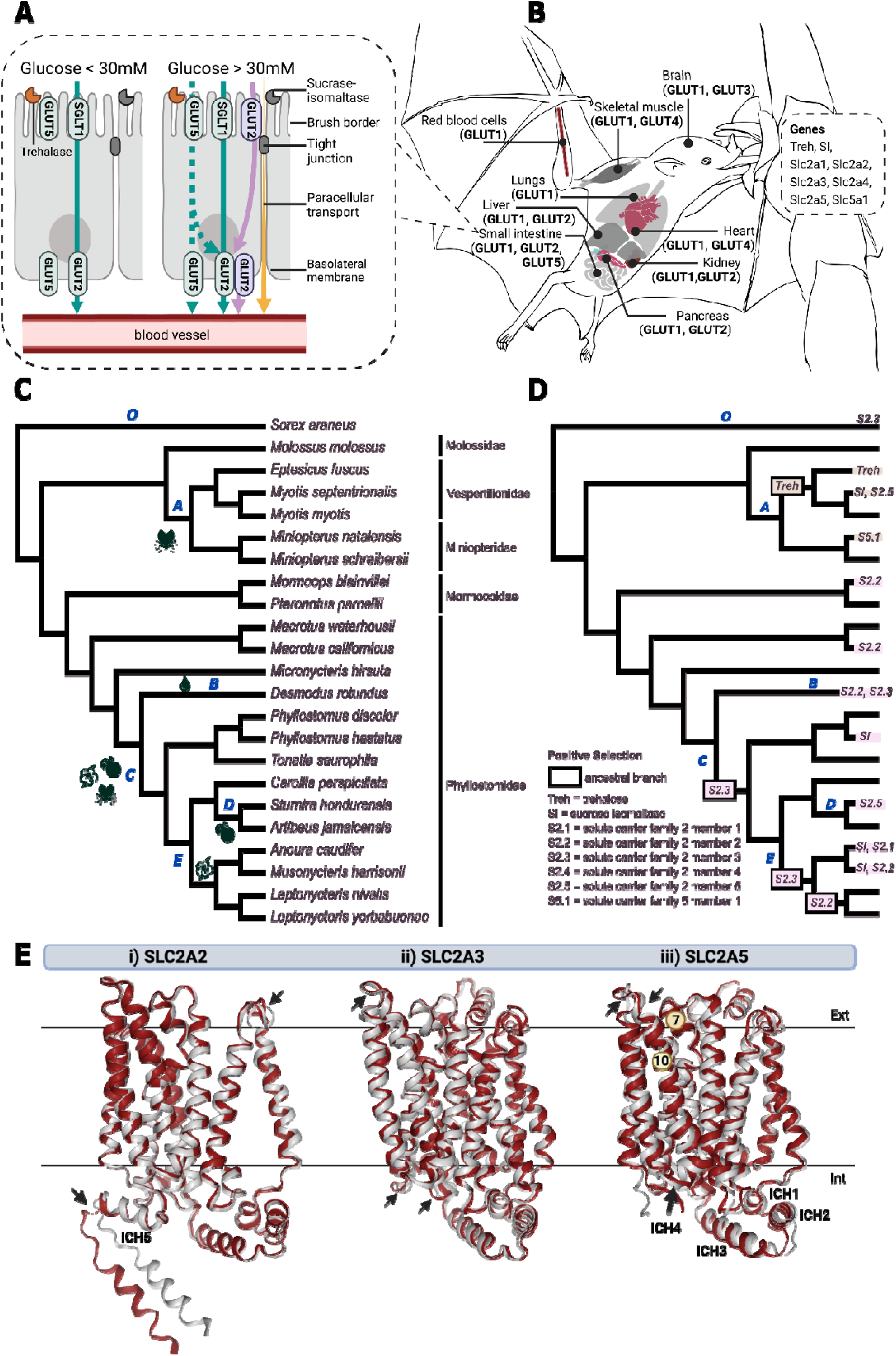
Molecular basis of sugar assimilation. A. Dietary sugar assimilation begins in the small intestine, along the brush border of enterocytes, where dietary enzymes trehalase and sucrase-isomaltase are located. Glucose transporters (SLC2A2/GLUT2, SLC2A5/GLUT5), sodium-glucose cotransporter (SGLT1), and paracellular transport determine the rate of glucose absorption into the bloodstream under low (<30mM) and high (>30mM) glucose concentrations. While GLUT5 is primarily a fructose transporter, it has the capacity to transport glucose (Burant et al. 1992, Mizuno et al. 2021). B. Glucose transporters move glucose from the bloodstream into specific tissues. Genes: Trehalase (*Treh*), Sucrase-Isomaltase (*SI*), Solute Carrier Family 2 Member 1 (*Slc2a1*), Solute Carrier Family 2 Member 2 (*Slc2a2*), Solute Carrier Family 2 Member 3 (*Slc2a3*), Solute Carrier Family 2 Member 4 (*Slc2a4*), Solute Carrier Family 2 Member 5 (*Slc2a5*), and Solute Carrier Family 5 Member 1 (*Slc5a1*). C. Species topology and foraging data follows Rojas et al. 2018. Insect-feeding branch leading to vesper and Miniopterus bats (A), blood-feeding (B), ancestral branch towards omnivores (C), fruit-eating (D), and nectarivore ancestral branch (E). D. Positive selection (p<0.01) of duodenal-enriched genes across ancestral and extant Neotropical bats with shrew as an outgroup (O). E. Ribbon representation of proteins, viewed in the plane of the membrane, from positive selected genes in the focal branch (Phyllostomidae) to outgroup (Mormoopidae). Foldseek structural comparisons of i) SLC2A2/GLUT2 from nectar bat *Musonycteris harrisoni* (gray) and insect bat *Pteronotus parnellii* (red); ii) SLC2A3/GLUT3 from nectar bat *Anoura caudifer* (gray) and *P. parnellii* (red); iii) SLC2A5/GLUT5 from fruit bat *Sturnira hondurensis* (gray) to insect bat *Mormoops blainvillei* (red). For each transporter shown, the C-terminal transmembrane (TM) bundle is located on the left and the N-terminal TM bundle is located on the right. The C- and N- terminal TM bundles transport glucose with a rocker-switch-type movement. TM7 and TM10 also support a gated-pore mechanism for glucose binding and release. The intracellular helices (ICH) domain provides stabilization for conformational changes. Arrows highlight structural changes where glucose binds extracellularly (ext) and where structural changes might affect substrate transport (int).

Using aBSREL (adaptive branch-site random effects likelihood; Smith et al., 2015) with HyPhy (Pond et al., 2005; 2020), we performed unbiased branch-site tests to identify episodic positive selection on orthologous gene sequences (Supplemental Table 4, 5). While positive selection implies that the genetic alterations within that gene have been advantageous, it remains challenging to relate selection signatures to functional adaptation. In contrast, proteins possess distinct three-dimensional configurations intricately associated with their functions. These structures generally undergo evolutionary changes at a slower pace than the sequences of DNA and amino acids constituting the proteins, given their close connection to protein function. Therefore, we created protein models for genes with signatures of positive selection using Alphafold (Jumper et al. 2021) and compared focal and outgroup species protein structure with Foldseek (van Kempen et al. 2023) to explore potential functional alterations.

Along the ancestral branch of the vespertilionid family (Fig.2C, D Branch A) and in the *Eptesicus* branch, comprising insectivorous bats, we detected signatures of selection in the trehalase gene (*Treh*). Surprisingly, the digestive enzyme sucrase-isomaltase gene (*SI)* and the transporter gene *Slc2a5* showed signs of selection in the vespertilionid lineage *Myotis.* To examine if genetic alterations might explain their enhanced ability to assimilate sucrose (i.e. elevated blood glucose levels), we examined the SI and SLC2A5 protein structure of two *Myotis* species and one *Eptesicus* species (Supplemental Figure 1). In line with our positive selection results, we found that SI within *Myotis* had TM-scorse of 0.7701 to 0.89313, whereas *Myotis* and *Eptiscus* had a TM-score of 0.9026. The greater differences in protein structure between *Myotis* species may reflect the ability of some species to tolerate sucrose. A common symptom of sucrose intolerance in insect-feeding bats (field observations), and in humans with sucrase deficiency, is intestinal distress from undigested sugar. A critical component of SI activity is its attachment to the plasma membrane of the gastrointestinal mucosa, facilitated by the P-type 1 domain of the isomaltase subunit (Hoffman and Hauser, 1993), which is one of the locations we observe differences in the *Myotis* SI structure (Supp Fig. 1B).

A single amino acid change in the second helical domain of the isomaltase subunit is sufficient to cause dissociation of SI from the membrane and the inability to process sugar (Jacob et al., 2000). It is possible that the changes in *Myotis*, compared to other insect-feeding bats like *Eptesicus,* provides the ability to process sugar, preventing them from having intestinal malabsorption. Variation is also predicted to be along the alpha and beta strands of both subunits, which have been known to cause dissociation of SI from the membrane and/or accumulation in the golgi (Jacob et al., 2000). For SLC2A5, the differences are located along the intracellular helices (ICH) domain, which anchor the conformational changes needed to transport glucose/fructose into the cell (Quistgaard et al. 2013). Similar missense substitutions to the ICH2-ICH3 domain of SLC2A5 are found in colon cancer cells (Tate et al. 2019), which are known to increase their glucose uptake (Warburg effect). Possibly, these modeled structural changes to the *Myotis* SLC2A5 similarly increase their glucose uptake (Supp Fig 1A).

In the vampire bats (Fig. 2C, Branch B), the ancestral branch leading to plant-eating (Fig. 2C Branch C), and in the ancestral branch leading to nectar-eating (Fig. 2C, node E), we found positive selection in the neuron-enriched *Slc2a3* gene (Fig. 2D). When comparing structural changes in SLC2A3 between the nectar bat *Anoura caudifer* and *Pteronotus parnellii* (Fig. 2E, ii), we obtained a TM-score of 0.97874. The amino acid substitutions are located along extracellular C-terminal transmembrane (TM) 10, where glucose binds, as well as within the plasma membrane, where it interacts with TM7 to coordinate their rocker-switch conformational changes with the N-terminal TM bundles during glucose transport (Deng et al. 2015). We also identified changes at the intracellular side of the C-terminal domain, possibly affecting the degree of opening during glucose transport. The TM-score of 0.9905 between *A. caudifer* and *Desmodus rotundus*, suggests that the transport of glucose into neurons of nectar bats and vampire bats is similar through this transporter. Possibly, these structural changes reflect more efficient glucose transport to meet the brain demands of these starvation-sensitive species (McNab, 1973; Freitas et al., 2013; Amaral et al., 2019).

The ancestral branch leading to the nectar genus *Leptonycteris* and the nectar bat *Musonycteris harrisonii*, as well as the lineage of vampire bats show signatures of selection in the gene *Slc2a2* (Fig. 2D, Branch B, E). The distinctive locomotor activities of these bats may subject them to similar selective pressures in meeting their increased energy demands of muscles, which extends to the brain (Bequet et al. 2001). Nectar bats have unique hovering flight capabilities (C. C. Voigt & Winter, 1999), while vampire bats are renowned for their proficiency in running (Ripperger & Carter, 2021). Increases in physical activity might increase the production of neurotransmitters, which are essential for brain function, and require glucose for energy. *Slc2a2* is also expressed in the gut and is largely responsible for affecting blood glucose levels (Ait-Omar et al., 2011), allowing glucose to be absorbed by brain tissue through the blood-brain barrier (Gerhart et al., 1992). It is possible that the signatures of selection on *Slc2a2* might reveal functional adaptations related to the ability of nectar bats to maintain high blood glucose at rest and the ability for vampire bats to assimilate glucose (Fig. 1A). When comparing predicted structural changes of SLC2A2/GLUT2 between *M. harrisonii* and *Pteronotus parnellii* (Fig. 2E, i), the TM-score was 0.9503. The differences are located along the extracellular N-terminal between TM bundles 4 and 5, which is usually highly conserved among sugar transporters (Nomura et al, 2015). *Slc2a2* is also expressed in the pancreas. We also predicted differences along the C-terminal end of the ICH5 domain, which undergoes post-translational modification for stability along the cell surface of pancreatic beta cells (Huttlin et al., 2010). It might be possible, that the stability of glucose sensing by the pancreas is transient in nectar and vampire bats, which might explain why these species have low levels of circulating insulin in response to glucose (Freitas et al., 2013; Castro et al., 2021).

In the ancestral branch leading to species with fruit diets, we observed no significant molecular adaptations, similar to a recent study on ancestral molecular evolution in Neotropical bats (Potter et al., 2021). However, we did identify a potential molecular adaptation related to glucose and fructose absorption in the genus *Sturnira* (Fig. 2C, Branch D). We detected positive selection for the gene *Slc2a5* and we found structural changes in SLC2A5, along the extracellular C-terminal domain where fructose/glucose binds (Fig. 2E, iii, arrows). We also found structural changes along TM10, which is known to work with TM7 to support a gated-pore mechanism for fructose/glucose transport. These are at least two structural changes in regions known to affect nutrient transport that might explain the distinctive *in vivo* blood glucose levels observed in *Sturnira* after sucrose ingestion (368 mg/dL), a pattern not observed in other fruit bats like *Artibeus* or *Dermanura* (Figure 1A).

In nectar and fruit bats (Figure 1A), we observed an increase in blood glucose levels 10 minutes after consuming sucrose, indicating the activity of the SI enzyme. Others have previously reported elevated sucrase activity in the intestines of nectar and fruit bats, in support of our observations (Schondube et al. 2001). While the nectar-feeding omnivore *Phyllostomus* (Figure 2C, branch C) did not assimilate significant amounts of glucose from digested sucrose (Figure 1A), they have the potential for elevated sucrase activity (Schondube et al. 2001). This suggests that SI may facilitate dietary expansion to omnivory in *Phyllostomus*, allowing them to respond to seasonal changes in food availability, which is in line with the fact that ingesting high-sugar sources, like nectar, is expected to increase sugar metabolism proteins (Kishi et al., 1999; Cui et al., 2003), particularly SI expression (Yasutake et al., 1995). Moreover, we found positive selection on the *SI* gene (Fig. 2D) in *Phyllostomus*, as well as in the nectar bats *M. harrisonii* and *A. caudifer,* accompanied by structural changes to the sucrase domain of SI (Supplemental Figure 1B). Detailed results of the aBSREL analysis are provided in Supplemental Tables 4 & 5, a summary of Foldseek results and the detailed Foldseek results are available from http://www.stowers.org/research/publications/LIBPB-2406.

### Intestine anatomy

We next investigated key anatomical features of the small intestine, with a focus on the duodenum. The first portion of the small intestine is where the majority of glucose absorption occurs, and the amount of absorbed glucose decreases as it progresses further along the intestinal tract (Fisher and Parsons, 1950; Riesenfeld et al., 1980; Wright et al., 2018). Consequently, the absorption through the duodenum represents the immediate pathway that glucose follows to reach the blood before reaching the jejunum or ileum. Since glucose assimilation did not show a phylogenetic signal, the traits we find in the intestine are more likely to be related to diet than to evolutionary relationships. In comparison to humans, bats lack circular folds in their small intestine (Silva et al., 2020), a finding that was confirmed in our analysis. We also observed striking differences in the intestinal length among bat species (Fig. 3), consistent with the study and drawings by Park & Hall (1951). By comparing bats with different diets, we found a correlation between the duodenum length and the proportion of sugar in their diet. Bats with rich-sugar diets, such as nectar and fruit bats, and also the insect-eating short-tailed fruit bat *Carollia* (York & Billings, 2009), exhibited longer duodenum, suggesting anatomical adaptations to deal with increased sugars and plant material, like fiber, as they shifted from insectivory to omnivory/frugivory. However, nectar bats do not consume high fiber levels, increased amino acids, or increased fatty acid quantities through their diets (Nicolson, 2022). Their diet mainly consists of water and a high concentration of sugar. Thus, we posit that simple sugars play a significant role in the shift in duodenum length.

**Fig. 3.**
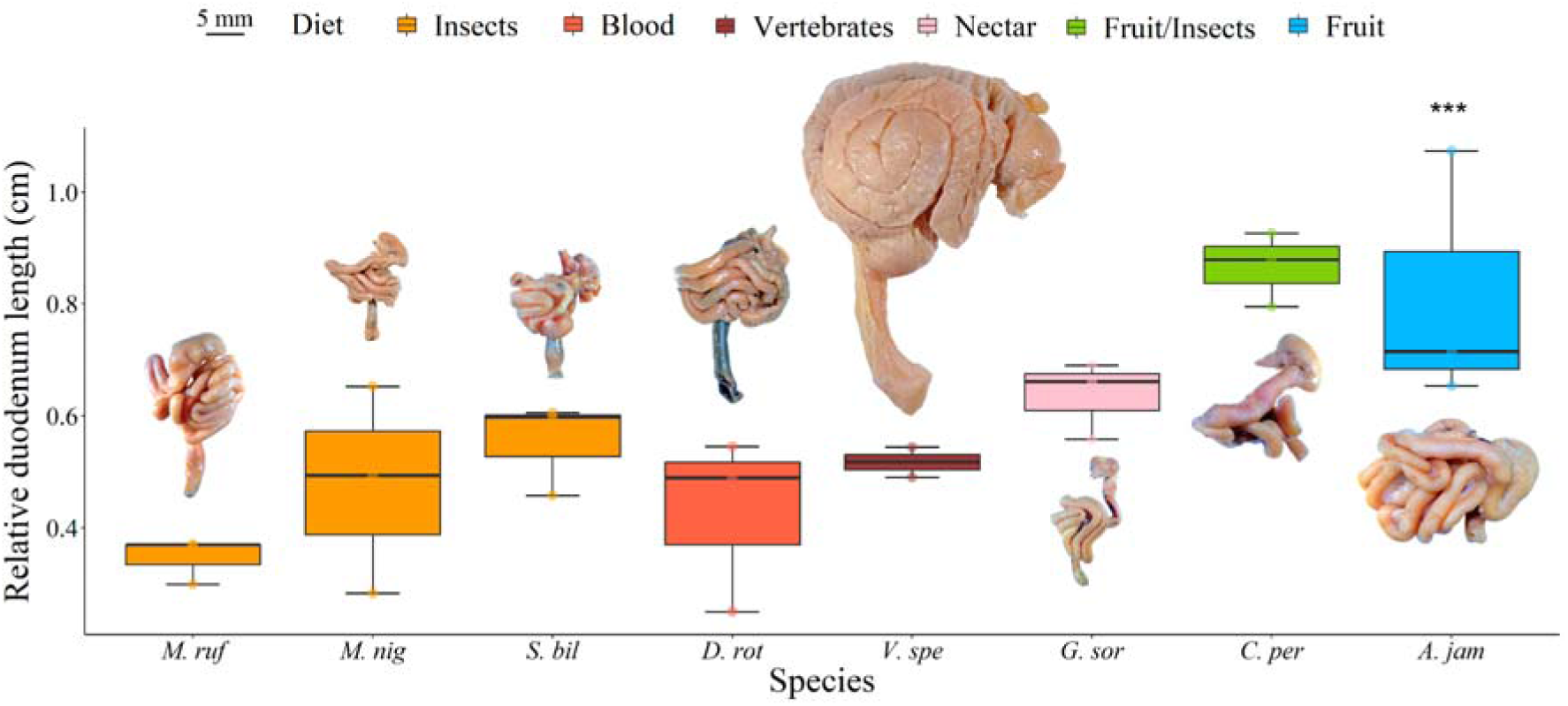
Relative length of duodenum associated with gastrointestinal tract morphology from eight bat species with different diets. Duodenum length relative to torso length in each species (shoulders to rump). *S. bil*: *Saccopteryx bilineata,* insectivorous; *M. nig: Myotis nigricans*, insectivorous*; M. ruf*: *Molossus rufus*, insectivorous; *D. rot: Desmodus rotundus*, hematophagous; *V. spe*: carnivorous; *C. per*: *Carollia perspicillata*, omnivorous; *A. jam*: *Artibeus jamaicensis*, frugivorous; *G. sor*: *Glossophaga soricina,* nectar-eating bat. Three individuals per species were reported, except *V. spe* with two individuals. Species with fruit and nectar diets tend to have longer duodenum than the blood, meat, and insect-eating species.

The intestine plays a crucial role in the absorption of nutrients, including sugars, amino acids, fatty acids, and vitamins, through specialized structures called villi that project into the lumen of the gut (Hilton, 1902; Walton et al., 2018). These villi consist of specialized epithelial cells (enterocytes) and goblet cells (Strobel et al., 2015), and in bats, they exhibit diverse shapes (Figure 4). Among the seven bat species examined for histology, we observed common finger-like villi (Fig. 4D #1 & #2), previously described pyramidal villi (Figure 4D #4), and zigzag villi (Figure 4D #6) (Scillitani et al., 2007; Gadelha-Alves et al., 2008; Silva et al., 2020). Additionally, we found villi shapes (Figure 4D #3 & #5) that, to our knowledge, have not been reported before. Across the duodenum and beginning of the jejunum, insectivorous bats exhibited finger-like villi, some of which had simple entrances along the border (Fig. 4D black triangles). *Glossophaga* exhibited entrances and sometimes an additional “gap” in the middle of the finger-like villi body (Fig. 4D #3 blue triangle). Pyramidal-shaped villi (Fig. 4D #4) were found in *Carollia*, *Artibeus,* and some *Myotis* individuals, while zig-zag villi (Fig. 4D #6) were mainly found in *Artibeus*, *Carollia*, with rare occurrences in *Myotis*. *Glossophaga* also displayed pyramidal and zig-zag villi but with more pronounced entrances along the entire length of the villi. We identified a novel villi type (Fig, 4D #5), that we referred to as “ruffled”, in *Carollia* and *Myotis,* characterized by multiple crests and troughs along its perimeter. The villi were scaled to the same size and their perimeter was measured. We found that while finger-like villi showed a perimeter of 339.56 µm, zig-zag villi were 931.99 µm, and pyramidal villi with a gap in the middle were 1124.73 µm. The diverse villi morphologies observed suggest functional adaptations aimed at increasing the absorptive area in bats’ intestine.

**Fig. 4:**
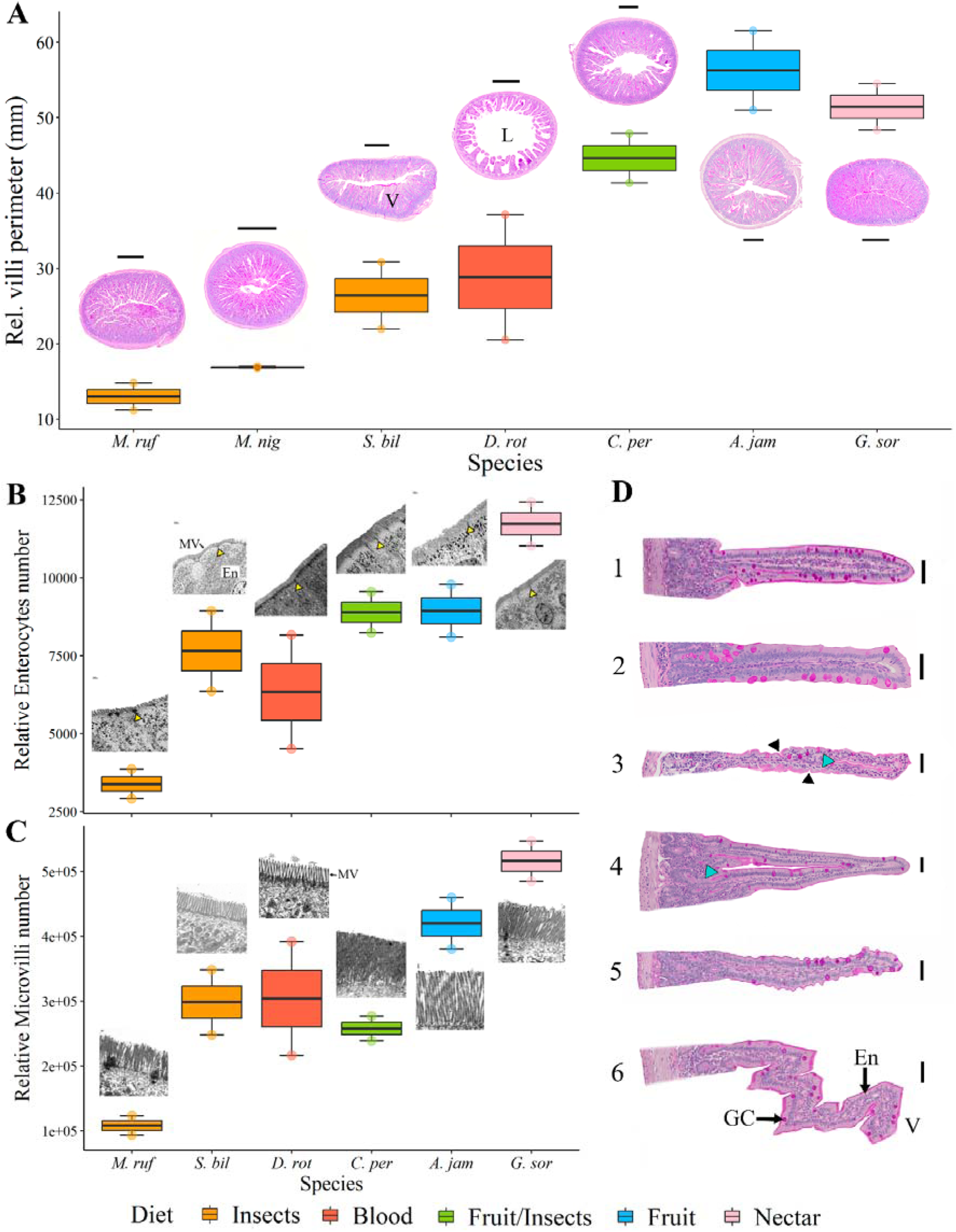
Histology of duodenum of bats with different diets. A. Relative villi perimeter across the duodenum related to cross-sections for each species; B. Relative number of enterocytes along the duodenum related to TEM of the enterocytes and microvilli; C. Relative number of microvilli along the duodenum related to TEM of microvilli. D. Different types of villi in bats, 1-2) finger-like villi, 3) finger-like villi with entrances (black triangle) and gap (blue triangle), 4) arch-like villi, 5) New villi “ruffled”; 6) zig-zag villi; (1, 4, 5 extracted from *C. per*; 2 from *S. bil*, 3 from *G. sor*, 6 from *A. jam*). Species *M. ruf*: *Molossus rufus, S. bil*: *Saccopteryx bilineata,*, *D. rot: Desmodus rotundus, C. per*: *Carollia perspicillata, A. jam*: *Artibeus jamaicensis*, *G. sor*: *Glossophaga soricina*. A and D. Periodic acid-Schiff-staining. Scale bars: A. 250 µm, D. #1-3, 5: 20 µm, #4, 6: 25 µm. Yellow triangles in B pointing at tight junctions between enterocytes (En). MV: microvilli. V: villi, L: lumen, GC: Goblet cells. Measurements were taken in different sections of the duodenum and are reported from one individual per species for the histology section and one individual per species for the TEM section.

Interestingly, the insectivorous *Myotis* exhibited higher sucrose assimilation compared to other insectivorous bats. We found evidence of positive selection for genes encoding SI and SLC2A5 as well as predicted changes to protein structure in the *Myotis* lineage. Additionally, we observed similar villi shapes in fruit, nectar, and omnivorous bats. *Myotis* was traditionally considered a strict insectivore; however, Novaes et al. (2015) reported fruit consumption in a species of this genus. This finding, along with predation on insects that feed on nectar (Mooseman et al., 2007), may be linked to similar trait evolution observed in bats with sugar-rich diets. Further ecophysiological investigations are necessary within this diverse genus and within phyllostomidae insectivorous species who might incorporate some plant material in their diets.

After villi shape analysis, we quantified the villus perimeter exposed to the lumen in the duodenum of species with different food preferences to assess its association with nutrient absorption. After adjusting for body size, we found a greater exposed surface area of villi in the duodenum for bats with sugar-rich diets (*Artibeus*, *Glossophaga*, and *Carollia*), compared to insectivorous and vampire bats (Fig. 4A). The variation in duodenum villi exposure, combined with the *in vivo* physiology data, suggests that the increased absorptive area in the duodenum of bats with sugar-rich diets may contribute to enhanced glucose absorption (Fig. 1A). A similar pattern has been observed in the proximal intestine of the distantly related frugivorous megabat *Epomophorus wahlbergi* (Makanya et al., 1997).

Using Transmission Electron Microscopy (TEM), we visualized the enterocytes and their microvilli, which contribute to a 20-fold increase in the absorptive surface area (Helander & Fändriks, 2014; Price et al., 2015). We measured individual enterocytes, defined as the distance between tight junctions, and quantified the number of microvilli per enterocyte. Our subcellular analysis revealed a higher number of enterocytes (Fig. 4B) and microvilli (Fig. 4C) in the corrected length of the duodenum of bats with rich-sugar diets, especially *Glossophaga* and *Artibeus* (maximum *in vivo* glucose levels above 600 mg/dL), compared to the insect and vampire bats, but especially compared to *Molossus* (maximum *in vivo* glucose levels of 174 mg/dL). The microvilli pattern observed in Neotropical bats aligns with discoveries in Old World bats, despite their divergence in the early Eocene. Old World insect-eaters have smaller and fewer microvilli, while Old World fruit bats have longer and more abundant microvilli (Makanya et al., 1997; 2005). The increased number of enterocytes and microvilli suggests enhanced paracellular absorption between intestinal cells (Caviedes-Vidal et al., 2008; Brun et al., 2014; Brun et al. 2019; Fasulo, 2014) and provides greater space for glucose transporters and digestive enzymes along the brush border (Price et al., 2015). These results indicate that bats have varying intestinal absorption capabilities, which broadly relate to diet. For example, in nectar and fruit-eating bats, the numerous microvilli could increase the density of SI and apical transporters (SLC5A1, SLC2A2, and SLC2A5), thereby promoting glucose absorption that leads to elevated blood glucose levels (Fig. 1). This hypothesis would fit with observations made on the increased levels of the sucrase enzyme expression in nectar and fruit-eating bats (Schondube et al. 2001).

### Molecular mechanism of glucose absorption by transporters (1461 words)

Next, we investigated the molecular activity underlying glucose transcellular absorption and its impact on plasma glucose levels. We examined the *in vivo* glucose transfer from the gut lumen into the enterocyte through the key components of molecular glucose transcellular absorption: Na^+^-dependent glucose co-transporter, SGLT1 (*Slc5a1* gene), the apical solute carrier family member 2 glucose transporter, SLC2A2/GLUT2 (*Slc2a2* gene), and SLC2A5/GLUT5 (*Slc2a5* gene) (Fig. 5) across seven bat species. The postprandial regulation of SI is not directly stimulated by glucose, and we confirmed this with qPCR (Supp Fig. 2), so we did not investigate these further. In non-insect feeding bats, a functional trehalase enzyme is absent due to inactivating mutations in the gene (Jiao et al. 2019), thus preventing further examination.

**Fig. 5:**
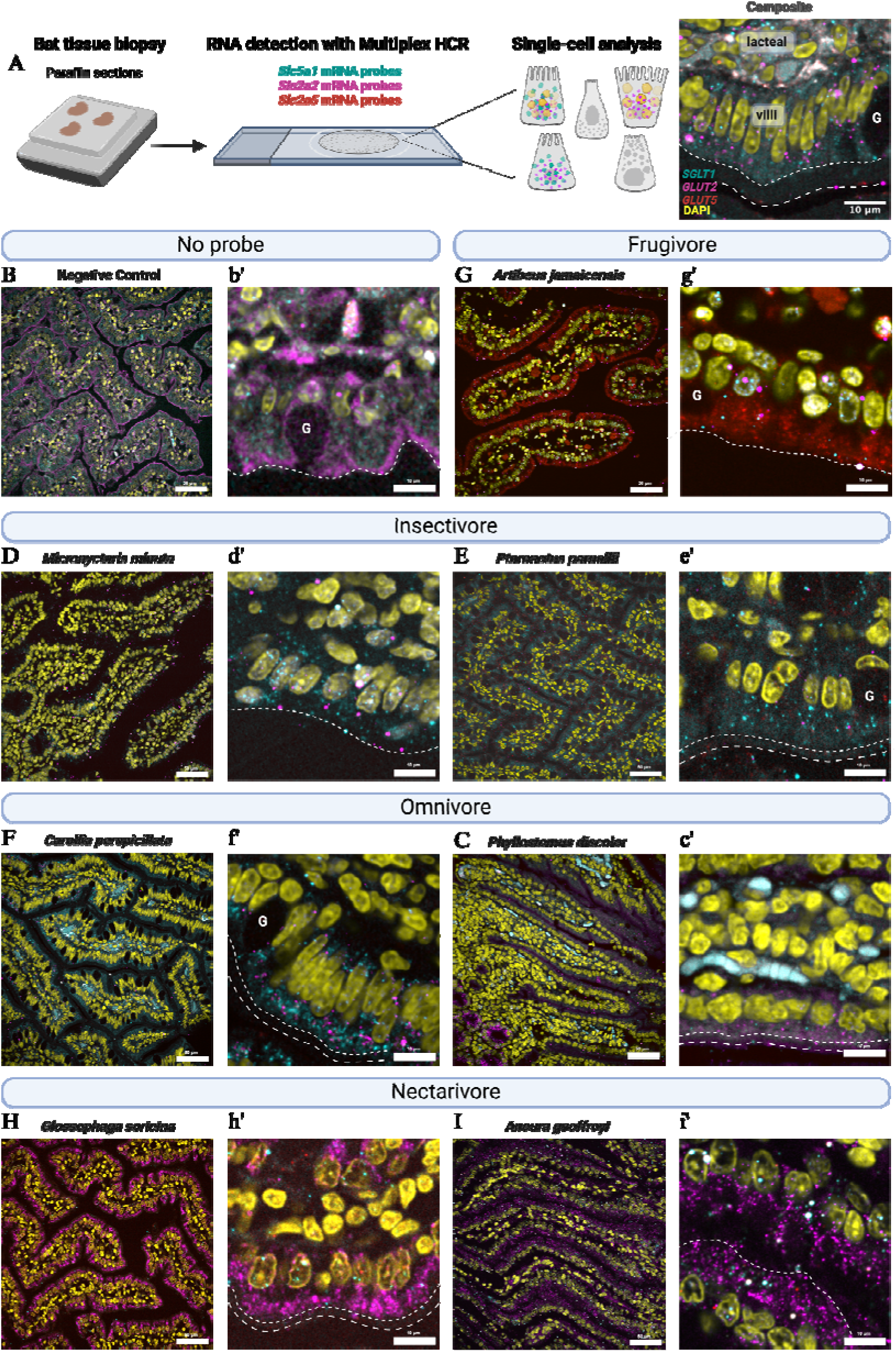
HCR RNA-FISH in the duodenum of bats from different diets. RNA expression is shown for bats fed a single dose (5.4mg/kg body weight) of glucose. A) We performed HCR RNA-FISH with probe sets targeting genes *Slc5a1*, *Slc2a2,* and *Slc2a5* on fixed paraffin-embedded bat intestinal tissue and quantified the RNA fluorescent signal per enterocyte. An example composite image of the duodenum with all probes labeled with anatomical features is shown. Apical epithelial layer shown with a fine-dotted line; microvilli brush border shown with a dashed line; goblet cells are denoted by “G”. B-I) 40x overview images of the intestinal villi for two omnivorous, two insectivorous, one frugivorous, and two nectarivorous species. B) Negative control with no probes, C) fruit; *Artibeus jamaicensis,* D) insect; *Micronycteris minuta\** and E) *Pteronotus parnellii,* F) omnivore; *Carollia perspicillata* and G) *Phyllostomus discolor* H) nectar; *Glossophaga soricina* and I) *Anoura geoffroyi*) at t=10 or *t=60. Scale = 50µm. b’-i’) Enlarged view of enterocytes along the villi. Scale = 10 µm. HCR probe sets are shown in cyan (*Slc5a1*), magenta (*Slc2a2*), and red (*Slc2a5*). DAPI (yellow) labels the cell nuclei. Bright, uniform, and large circular spots (white) are background noise from amplifiers.

We employed multiplexed, single-molecule HCR™ RNA-fluorescent in situ hybridization (RNA-FISH) on fasted bats (t=0) and bats that were fed (t=10) a single dose of glucose (5.4g/kg body weight) with a 20% glucose solution (∼134mM of luminal glucose, assuming complete gastric emptying from the pylorus). We investigated two species within each dietary category (Fig. 5): *Pteronotus parnellii* (n=4) and *Micronycteris minuta* (n=1) for insect-eating bats, *Carollia perspicillata* (n=4) and *Artibeus jamaicensis* (n=2) for fruit-eating bats, and *Glossophaga soricina* (n=4) and *Anoura geoffroyi* (n=4) for nectar-feeding bats. Additionally, we examined one omnivorous bat species, *Phyllostomus discolor* (n=1). We observed a notable pattern where nectar bats consistently displayed a high *Slc2a2* signal in both fasting and fed states, whereas the fruit bat *A. jamaicensis* exhibited the highest *Slc2a5* signal (Fig. 5). The fruit bat *C. perspicillata,* which includes a significant amount of insects in its diet, exhibits similar gene expression as the insect bats and the omnivore *P. discolor.* This supports *Carollia* classification as an omnivore (York & Billings 2009) with a large proportion of fruit in their diet.

Next, we quantified the fluorescent signal intensity of each HCR probe within a single enterocyte along the mid-length of the villi (Fig. 6). Additionally, we performed RT-qPCR to observe broad patterns in RNA expression against the housekeeping gene GAPDH (Supp Fig. 2). In response to dietary glucose, we observed a pattern of response similar to that described in mouse studies (Shirazi-Beechey et al. 2011), although the extent of this response varies among species (Supp Fig. 4). In the insectivorous bat, *Pteronotus parnellii*, enterocytes responded with a log2 fold change of 1.8 in *Slc5a1* expression, 1.1 in *Slc2a2* expression, and 2.6 in *Slc2a5* expression, accompanied by a rise of 148 mg/dL in their blood glucose levels (Supp Table 6). Comparatively, the insect bat *Micronycteris minuta*, had lower expression for *Slc5a1*, similar expression of *Slc2a2,* and no detected expression of *Slc2a5* in response to glucose (Fig. 6, Supp Fig. 4C), with a change of 174.7mg/dL in blood glucose (Supp Table 6), suggesting the importance of *Slc2a2* with rising blood glucose levels. On the other hand, the individuals tested here from the species *P. discolor*, (omnivorous) elevated blood glucose levels to 500 mg/dL. However, despite the absence of a time-series, the expression of *Slc5a1* and *Slc2a2* in response to glucose in this species did not differ significantly from that in *P. parnelli* or *C. perspicillata* (Supp Fig. 4). The insect-eating, short-tailed fruit bat *C. perspicillata*, exhibited an increase of 440 mg/dL in blood glucose levels (Supp Table 6), a log2 fold change of 1.1 in *Slc5a1* expression, 0.468 in *Slc2a2* expression, and 1.11 in *Slc2a5,* much lower than the response of *P. parnellii*. RT-qPCR data further highlights that *P. parnellii* exhibits broader change in glucose transporter activity in response to glucose, compared to more specialized phyllostomids (Supp Fig. 2). This data suggests that the dramatic rise in blood glucose levels in *C. perspicillata* and *P. discolor* can be primarily attributed to the paracellular absorption, while in the more specialized groups the transporters’ gene expression changes are more specific (Fig. 6, Supp Fig. 2).

**Fig. 6.**
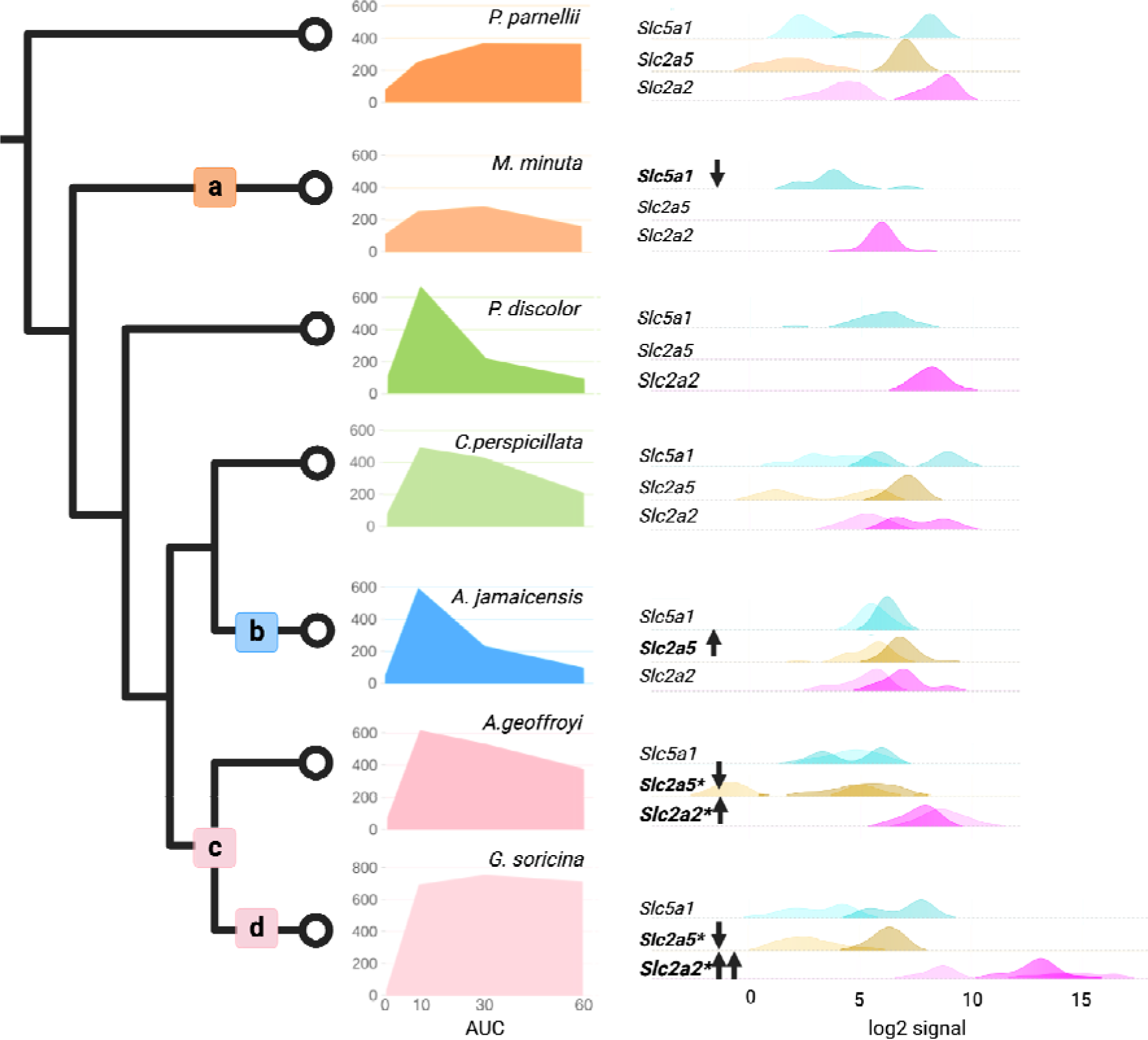
Parsimony-based evolutionary inferences of blood glucose levels in relation to gene expression. Average species blood glucose levels across 60 minutes (AUC, area under the curve) and change in gene expression 10-minutes after eating a 20% glucose solution. Most dietary guilds are represented by two species: insect, *P. parnelli, M. minuta*; fruit, *A. jamaicensis*; nectar, *A. geoffroyi*, *G. soricina*. *P. discolor* and *C. perspicillata*, are omnivores incorporating large proportions of nectar or fruits, respectively, in their diet. Right panel, results of the single-cell image analysis taken from a minimum of 20 cells at fasted (t=0) and fed (t=10), presented as ridgeline density plots colored by HCR probe set as follows: *Slc5a1* (cyan), *Slc2a5* (yellow), *Slc2a2* (magenta). The log2 values of fluorescent intensity are displayed for t=0 in a lighter shade and t=10 in a darker shade. Shared gene expression responses (no significant differences) are taken as evidence for inheritance from a common ancestor. Significant gene expression differences among species (p<0.001) are noted as apomorphies on the tree as follows: a) a decrease in *Slc5a1* in *M. minuta* and a decrease in blood glucose levels (200mg/dL), b) an increase in *Slc2a5* expression in *A. jamaicensis* and an increase in blood glucose levels (600mg/dL), c) a decrease in *Slc2a5* expression and an increase in *Slc2a2* gene expression in *A. geoffroyi* and *G. soricina*, with an increase in blood glucose (>600mg/dL), and d) the highest expression of *Slc2a2* and the highest reported blood glucose levels in *G. soricina* (>750mg/dL).

The fruit-eating bat *A. jamaicensis* increased blood glucose levels by 483mg/dL upon eating sugar, but only displayed a log2 fold change of 0.006 in *Slc5a1*, 0.350 in *Slc2a2*, and 0.294 in *Slc2a5*. While expression changes were minimal, the initial fasting levels of *Slc5a1* and *Slc2a5* were significantly higher than those in *P. parnellii* and *C. perspicillata* (Supp Fig. 4), indicating that gene regulatory changes, in addition to paracellular absorption, may explain some of the rapid increase of blood glucose levels at the first ten-minutes interval (Fig. 6). The nectar specialists, *A. geoffroyi* and *G. soricina,* showed the highest blood glucose change of 639 mg/dL and 621mg/dL, respectively (Supp Table 6). Using a more specialized glucometer (AlphaTRAK® 2.0), we obtained blood glucose readings of more than 750 mg/dL (Supp. Table 6). These levels are to our knowledge, the highest documented for any wild species feeding on a single sugar dose of glucose solution. These two bats showed significantly elevated expression in *Slc2a2* at fasting (Supp Fig. 2 & 4C) and the lowest log2 fold change in *Slc2a2*, −0.123 and 0.109, respectively (Supp Table 6). Both nectar bat species also had significantly lower expression of *Slc2a5* (Supp Fig.4B). These findings suggest the significance of regulatory changes in *Slc2a2* per enterocyte regarding the rapid rise in blood glucose levels.

The evolutionary patterns of gene expression demonstrate similar responses to glucose among bat species. When considering the broad patterns of blood glucose levels in relation to the absorption of glucose via the expression of intestinal transporters and increase in microvilli (Fig. 4C), we observe molecular apomorphies in transporter gene expression (Fig. 6). These differences may contribute to the diverse blood glucose patterns observed in the insectivore *M. minuta* (Fig. 6a), the frugivore *A. jamaicensis* (Fig. 6b), and the nectar bats (Fig. 6c,d). To relate how much and how long blood glucose levels remain elevated after the consumption of a glucose solution, we used a summary measure of the overall glucose response (Fig. 6, AUC). Relative to the outgroup *P. parnellii,* a higher or lower AUC indicates a change in glucose assimilation. For example, *Slc5a1* expression is lower in *M. minuta* (t=60) and we observe an AUC_30-60_ of 6705 (Fig. 6a), in contrast to *P. parnellii*, which has higher expression of *Slc5a1* and an AUC_30-60_ of 9630, indicating a diminished transfer of glucose through *Slc5a1* in *M. minuta* from the intestinal lumen to the bloodstream. Comparatively, the expression levels of transporters for *P. discolor* (AUC_0-10_ of 3312) and *C. perspicillata* (AUC_0-10_ of 2828) were similar to those of *P. parnellii* (AUC_0-10_ of 1368). Since the expression in each enterocyte is similar, the rise in blood glucose levels is likely the result from an increase in the number of enterocytes and increase in paracellular absorption, made possible by an elongation of the duodenum (Fig. 3), and an expansion of the villi perimeter (Fig. 4). On the contrary, *A. jamaicensis* has an increase in *Slc2a5* expression (Fig.6b) and an AUC_0-10_ of 2973. The increased expression of glucose transporters may contribute to the rise in blood glucose levels, given that *A. jamaicensis* exhibits similar enterocyte numbers, and thus potential paracellular absorption, as *C. perspicillata* (Fig. 4B). The regulatory changes of *Slc2a5* may be an apomorphic molecular change in the genus *Artibeus* (Fig. 6b) or it may be a plesiomorphic change in the subfamily Stenordermatidae, a specialized group of fruit-eating bats. It is worth mentioning that *Sturnira’s* molecular adaptations to SLC2A5 (Fig. 2) might have been facilitated by the consistent usage of this molecular transport system. While fruit-eating bats have increased *Slc2a5*, nectar bats have decreased it, instead depending extensively on *Slc2a2* to increase blood glucose levels (Fig. 6c, d). The changes in the timing of *Slc2a2* and the amount of *Slc2a2* and *Slc2a5* expression likely reflect molecular plesiomorphies in the nectar bat subfamily Glossophaginae (Fig.6c). A possible molecular apomorphy is represented by the excessively elevated levels of *Slc2a2* within the genus *Glossophaga* (Fig. 6d, Supp Fig. 4). Correspondingly, *A. geoffroyi* (AUC_0-10_ of 3842) and *G. soricina* (AUC_0-10_ of 3636) have extremely high blood glucose levels (average >700mg/dL; Supp Table 6) compared to other bats.

This study signifies the first case of persistent intestinal *Slc2a2* expression found in a non-pathologic state in mammals. A comparable observation has been reported in the Ruby-throated Hummingbird, providing a notable example of convergent evolution. (Ali et al., 2020). The increase in *Slc2a2* expression also exemplifies repeated evolution toward a diet with a high sugar proportion, a molecular trait observed in Old World fruit bats such as *Rousettus leschenaultia* and *Cynopterus sphinx*, which have maximum blood glucose levels between 430 and 490 mg/dL (Meng et al., 2016). These species, as well as Neotropical bats, such as *Carollia*, *Artibeus*, and *Anoura*, have an 11-base pair deletion in the proximal promoter of the *Slc2a2* gene. This deletion is predicted to interfere with transcriptional repressors to enhance gene activity. While we did not observe a significant increase in *Slc2a2* expression in *Carollia* and *Artibeus* at the level of the enterocyte, it might be altered in the liver cells, where *Slc2a2* is also expressed, and might explain their increased liver glycogen stores compared to nectar bats (Pinheiro et al. 2006; Amaral et al. 2018). Future studies on tissue-specific gene regulation are needed to uncover mechanisms of evolutionary change across species.

Given the high glucose concentration present in their liquid nectar diets and their extremely high postprandial glucose levels, results suggest that the distinctive expression pattern of *Slc2a2* is attributable to a specific adaptation of nectarivorous bats (Fig. 6c). The permanent presence of apical *Slc2a2* might result from defective insulin action, as insulin typically triggers the internalization of GLUT2 (*Slc2a2)* to slow down sugar uptake and prevent high blood glucose levels after a sugar meal (Tobin et al., 2008). This is in line with the fact that nectar-feeding bats need to absorb high quantities of glucose to fuel their energetic requirements (Suarez & Welch, 2017; Voigt & Speakman, 2007) and the hypothesis about additional mechanisms, complementary to insulin, to regulate glucose homeostasis (Kelm et al., 2011; Peng et al., 2017). The magnitude of postprandial hyperglycemia observed in nectar bats provides strong evidence that the adaptation towards nectar feeding has primed the duodenum for immediate and enhanced dietary sugar absorption, ensuring an adequate supply of glucose to tissues. The metabolic adaptation is further supported by the permanent apical expression of *Slc2a2* associated with obesity and diabetes, conditions characterized by elevated blood glucose levels, insulin resistance, and an increase in villous surface area (Marks et al., 2003; Ait-Omar et al., 2011; Gromova et al., 2021). Therefore, the long-term dietary adaptation to excess sugar consumption required the modification of the fundamental regulation of glucose absorption at the enterocyte level involving transporter trafficking in addition to the well-known paracellular absorption mechanism.

## Conclusion

Focusing on three dietary sugars—trehalose, sucrose, and glucose—we unveil a remarkable spectrum of adaptations within the glucose homeostasis pathway. In our investigation into the metabolic adaptations across more than 29 bat species with diverse diets, we found higher assimilation of glucose and sucrose in nectarivorous, frugivorous, and certain omnivorous bats, whereas insectivorous and omnivorous bats exhibited greater trehalose assimilation. Intriguingly, no insectivorous bat showed sugar absorption and assimilation as rapid and extensive as bats with sugar-rich diets. The observed variations in metabolic phenotypes are intricately linked to distinct adaptations in digestive morphology, including alterations in intestinal length, exposed villi, and microvilli. Bats with sugar-rich diets exhibited a longer duodenum and higher numbers of enterocytes and microvilli along the initial section of the small intestine. These features suggest an enhanced capability for glucose absorption in bats with sugar-rich diets. Moreover, our study identifies key genetic traits associated with efficiently extracting maximal glucose energy from dietary inputs. Positive selection is evident in genes encoding for sucrase-isomaltase (*SI*) and glucose transporters in nectar and omnivorous bats and on a fructose transporter in a fruit bat. Structural comparisons of these proteins further elucidate the impact of amino acid substitutions on their functional roles, which may change enzymatic reaction speed or the affinity of glucose to transporters. Notably, our investigation extends beyond genetic traits to explore shifts in gene expression within single enterocytes along the brush border of the duodenum. This detailed examination of transporters SLC2A2/GLUT2 (*Slc2a2*), SLC2A5/GLUT5 (*Slc2a5*), and SLC5A1/SGLT1 (*Slc5a1*) unveils a nuanced interplay in response to dietary sugars. Across most bat species examined, there is a conserved expression response of sugar assimilation genes to glucose per enterocyte, so gut morphology appears to be the primary driver for glucose assimilation differences. As bats must limit gut size due to the constraints imposed by flight, their guts’ microanatomy gets modified. It could also be that the structural changes we have documented make each transporter more efficient at its function while keeping gene expression the same. The continuous expression of *Slc2a2* encoding for glucose transporter GLUT2, was exclusive to nectar bats, indicative of enhanced glucose receptivity in this group. This interplay between physiology and ecology over evolutionary time illuminates the intricate adaptive mechanisms that underlie diet evolution in bats. The overarching narrative underscores the sophisticated adaptability of absorptive villi and highlights its relevance to conducting further digestive physiology studies, including those involving more taxa.

Our integrative approach reveals that the differences in sugar assimilation among Neotropical bats directly correlate with their dietary evolution and provide valuable insights into metabolic adaptations. Digestive enzymes, glucose transporters, and absorptive gut traits have undergone adaptive selection, enabling the transition from an insect-eating ancestor to a plethora of species with diversified diets. We have unraveled the intricately linked mechanisms underlying sugar assimilation in these remarkable mammals through a multidisciplinary approach, encompassing *in vivo* physiology measurements, genomics, gut anatomy, and single cell analysis. These findings have broader implications for our understanding of blood glucose regulation and provide a foundation for future investigations into the interplay between genotype, phenotype, and ecological factors. Further exploration of the molecular and physiological adaptations in bats will undoubtedly enrich our understanding of the evolutionary dynamics that shape the diversity of life on our planet.

## Methods

### Field work

The fieldwork was made under the permit of the National Authority of Environmental Licenses and the Ministry of Environment and Sustainable Development of Colombia, Resolution 1070, August 28^th^, 2015. Bats were captured between June 2019 and December 2020 in 11 localities of the dry tropical forest ecosystem in the department of Valle del Cauca, Colombia, some within the Dry Tropical Project from the Institute for Research and Preservation of the Cultural and Natural Heritage of Valle del Cauca (INCIVA). To catch species with different food preferences (frugivores, insectivores, nectarivores, omnivores, and hematophages), we opened mist nets between 18:00 and 24:00 h. In some instances, bats were manually captured in their refuge. Each individual was taxonomically identified, and their weight, age, sex, reproductive status, and diet type were recorded. We also include species captured in Trinidad and Tobago as well as Costa Rica. We acknowledge that the bat’s diet is more a continuum rather than static categories, however, for analyzing the data we grouped species according to their food preference and morphological adaptations for the consumption of the food resources (Park & Hall, 1951; Freeman, 1998; Rojas et al., 2011; Santana et al., 2011). In this way, the species of the genera *Artibeus*, *Dermanura*, *Uroderma* and *Sturnira* are considered frugivores, *Carollia* and *Phyllostomus* as omnivores, *Glossophaga*, *Choeroniscus* and *Lonchophylla* are considered nectarivores, *Desmodus* as hematophagous, *Vampyrum* as carnivore and *Saccopteryx*, *Peropteryx, Myotis* and *Molossus* and *Pteronotus* as insectivorous. We include data (for the *in vivo* physiology section) for 79 individuals with a preference for fruits, 55 omnivorous bats, 23 with a preference for nectar, 27 for insects, 14 for blood, and 2 carnivorous bats (Supplementary information, Table 1). Juvenile individuals and pregnant or lactating females were excluded from the study due to high energy requirements and significant physiological changes during these stages, compared to non-pregnant or non-lactating adult individuals (Kurta et al., 1989; C. Voigt, 2003).

### 1.2 Oral glucose tolerance tests

After identification, bats were fed a 20% sugar solution and subsequently subjected to a period of fasting for ten to twelve hours. After fasting, each bat was fed a bolus of sugar (5.4g/kg of body weight) as previously established by Kelm et al. (2011). Individual bats were only fed one type of sugar (Glucose, Sucrose, or Trehalose). To determine blood glucose levels, a drop of blood was drawn from the forearm with a 30G lancet before the sugar bolus and 10, 30, and 60 minutes post-feeding. The blood was immediately measured using a GlucoQuick G30a glucometer (Diabetrics®) with a 20-600 mg/dL range. The individuals corresponding to *Lonchophylla* and *Pteronotus* were captured in Costa Rica and Trinidad and Tobago in 2022, respectively. Their blood glucose levels were measured with an AlphaTRAK 2.0 glucometer (Zoetis®) with a range of 20-750 mg/dL, however the measurements above 600 mg/dL were treated as 600 measurements to match the rest of the data previously obtained with the more limited range glucometer.

The bats individually remained in cloth bags between readings. Finally, the bats were tagged, fed, and released. In total, blood glucose was measured for 199 individuals from 29 species and five families: Phyllostomidae, Mormoopidae, Emballonuridae, Vespertilionidae, and Molossidae.

#### 1.2.1 General patterns of sugar assimilation curves

We proposed general curves (Figure 1B) to describe the temporal pattern of assimilation of different sugars in Neotropical bats based on a 1-hour glucose tolerance test. The glucose curves were classified according to the speed of sugar assimilation to facilitate the understanding of the behavior of the curves. The *“Fast”* assimilation curve shows a peak of blood glucose levels only after 10 min of sugar ingestion, followed by a decrease in blood glucose levels. The *“Medium”* assimilation curve shows a peak of blood glucose levels 30 min after sugar ingestion followed by a decrease in blood glucose levels. The *“Slow”* assimilation curve shows a continuous increase in blood glucose levels until the final time point, with the maximum levels 60 min after sugar ingestion. Finally, the *“Limited”* assimilation curve shows blood glucose levels with little variation throughout the different time points of the test.

### 1.3 Statistical analysis

Blood glucose levels of all bats measured were grouped by species and plotted in R (Figure 1). Afterward, we evaluated the phylogenetic signal for the assimilation of the various sugars by the Pagel λ index (Pagel, 1999). We measured assimilation as the area under the glucose tolerance curve corrected (AUCc), this is, setting the base of the curve at the minimal blood glucose level recorded for the average measurements of each genera. Additionally, to test for the interactive effect of time points and food preferences on the assimilation of sugars, by considering various individuals per species and the non-independence due to shared evolutionary history among taxa, we implemented a Bayesian multilevel phylogenetic model in the brms R-package (Bürkner, 2018). This multilevel model includes a varying intercept over species (using an indicator variable for species) and a covariance matrix to specify the lack of phylogenetic independence, allowing the analysis of hierarchical biological data while incorporating evolutionary relationships among species. For statistical inference, we compared 95% credible intervals from the models between time points and food preference combinations. We calculated the phylogenetic correlations among species from an ultrametric tree using the vcv function of the ape R-package (Paradis & Schliep, 2018). For phylogenetic analyses, we used the species-level mammal phylogeny of Upham et al. (2019) (http://vertlife.org/phylosubsets/), so we downloaded a credible set of 10,000 trees for all taxa we have data on sugar assimilation and computed the consensus tree with the averageTree function of the phytools R-package (Revell, 2012). We performed the analysis using R 4.3.1 (R Core Team, 2023).

### 2.1 Morphology and histology of the GI tract

We extracted the GI tract from individuals preserved in 70% ethanol from species obtained in previous studies on bats (Camacho et al., 2020) obtained under the Wildlife Section, Forestry Division, Ministry of Agriculture, Land and Marine Resources (Republic of Trinidad and Tobago) permit number 1737 (2014). The species incorporated in this section were *Glossophaga soricina*, *Artibeus jamaicensis*, *Carollia perspicillata*, *Molossus rufus* and *Saccopteryx bilineata*. Additionally, we included two species obtained from the KU Biodiversity Institute and Natural History Museum: *Myotis nigricans* (KU 134846) and *Desmodus rotundus* (KU 100374), and one species obtained from the American Museum of Natural History: *Vampyrum spectrum* (M-267445, 272936). Finally we added more individuals to the analysis from the AMNH (*A. jamaicensis* 7473; *C. perspicillata* 7458, 175795; M. rufus 178675, 178672; *S. bilineata* 184693, 149982; *M. nigricans* 175725, 175724; *D. rotundus* 175406, 239943). We measured the length of the duodenum by imaging the unfolded-stretched intestine of each species with a Canon EOS Rebel E7i. Then, we determined the length through the Fiji platform – Image J 1.53f51 (Schindelin et al., 2012), and we compared relative intestine length as the gut length divided by torso length (shoulders to rump length). We used this measure to control by body size due to our focus on the intestine and its location, without adding measures related to the length of the rostrum or the tail, which can vary across bat species and families.

We investigated the duodenum because it is the main segment of the small intestine responsible for glucose absorption, and in vertebrates the amount of absorbed glucose decreases as it reaches the next intestinal sections (Riesenfeld et al., 1980; Fisher and Parsons, 1950). In this way, the duodenum was further analyzed with histology. The duodenum is the first section of the intestine from the end of the stomach-pylorus to the duodenojejunal flexure (Ishida et al., 2018). The duodenum was recognized by its greater diameter compared to the jejunum, the following intestine section after the duodenum. The jejunum was also investigated through histology, however its length could not be assessed due to the external similarity between the end of the jejunum and the start of the ileum.

For histological comparisons, we cut 0.5-1 cm sections of the duodenum. The gut tissues were embedded in paraffin, serially cross-sectioned to 10 µm, and stained with periodic acid– Schiff (PAS). All tissues were imaged using an Olympus slide scanner equipped with a 20X objective and exported as a TIFF. Two sections with good morphology per individual were chosen for study. We measured the area of villi and lumen (VLA) in each cross-section, and then we delimited 1/10 of VLA to measure the well-preserved villi perimeter (VPSA) in this sample area (SA). We also measured the lumen area (LA) to calculate the total villi perimeter along the cross-section. Then, we corrected the measurement for size differences among species by considering the cross-section perimeter (CSP) from the different species. Finally, we extrapolated the final value from villi exposed in the whole cross-section, corrected by size, to the relative duodenum (D) length (Eq. 1). (Measures summarized in Supplemental Figure 3).

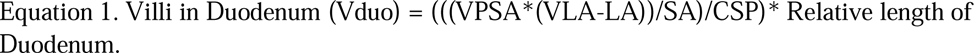

### 2.2 Transmission electron microscopy

To look closer at the enterocytes and microvilli, we used transmission electron microscopy (TEM). We sectioned 0.5-1.0 cm of duodenum from some of the species mentioned above (Fig 3). The samples were rehydrated and fixed in 50 mM Sodium Cacodylate (pH 7.4) containing 2.5% Paraformaldehyde and 2% Glutaraldehyde. The tissue segments were then post-fixed with 2% OsO4 for 2 hours, washed, and stained with 1% aqueous Uranyl Acetate overnight. After dehydration with a gradient of Ethanol, the samples were infiltrated and embedded into Epon resin. Ultrathin (80 nm) sections were cut with a Leica UC7 Ultramicrotome, collected onto slot copper grids, and stained with 4% Uranyl Acetate in 70% Methanol and Sato’s triple lead solution. Sections were imaged using a FEI transmission electron microscope at 80kV using the DigitalMicrograph® software.

To calculate the number of enterocytes in the duodenum we measured the enterocyte width of five different cells from each species, and the average was extrapolated to the relative length of the duodenum by dividing the villi surface found in the duodenum by the enterocyte width average. In addition, for calculating the number of microvilli in the duodenum, we counted the number of microvilli in each of the five different enterocytes where we measured the width, and then we multiplied the average microvilli value by the number of enterocytes. All the measurements were made with the Fiji platform – Image J 1.53f51 (Schindelin et al., 2012).

### 3.1 Genomics

Genome assemblies for 22 Chiroptera species, which included representatives from each of the dietary groups, were downloaded from online sources (Supplemental Table 1). Transcriptomes and proteomes for 16 species were obtained from the National Center for Biotechnology Information (NCBI) and Ensembl databases (Supplemental Table 1). After filtering out genomic contigs less than 10kb we generated gene models using MAKER (v3.01.03) using the 16 transcriptomes and proteomes (plus SwissProt/Uniprot v2021_03) as “EST evidence” and “Protein Homology Evidence”. RepeatMasker (open-4.0.7 with DB version 20170127) was used by MAKER to identify mammalian repeats. Links to the versions of all publicly available data (genome assemblies, transcriptomes, proteomes) used by MAKER can be accessed on SIMRbase (https://simrbase.stowers.org/bats_sugar_assimilation). Our SIMR bat models were assigned names derived from Uniprot (UniProt consortium, 2020) best hit with an e-value <= 1e-5. SIMR bat models are available for download and for browsing on SIMRbase (https://simrbase.stowers.org/bats/data). Genome browsers were built using Jbrowse (Buels et al., 2016) and SIMR buelsbase was built using Tripal (Spoor et al., 2019).

### 3.2 Ortholog Assignment and Candidate Gene Identification

To relate the blood glucose tolerance assay to evolution, we identified genes involved in sugar assimilation, specifically from the gut absorption into the bloodstream. A short list of eight genes (Table S3) were identified as having biased function and expression in the duodenum using mouse ENCODE (NCBI Bioproject PRJNA66167) and human HPA (NCBI Bioproject PRJEB4337). Orthologs were assigned to the SIMR bat models with standalone Orthologous Matrix (OMA) (version 2.4.2) using three species included in the OMA FASTA database (April 2021), *Homo sapiens* (HUMAN), *Pteropus vampyrus* (PTEVA), and *Myotis lucifugus* (MYOLU). A gene is considered an ortholog if it is found in the same OMA ortholog group (OG) as a HUMAN candidate gene with appropriate HGNC identifiers (Table S4).

### 3.3 Phylogenetic analysis of sugar related genes

Whole genome alignments of single-copy orthologs trimmed with trimAl v1.4.rev15 (Capella-Gutiérrez et al., 2009) from 22 SIMR bat models and two outgroups (Supp Table 1) were used to generate a phylogenomic tree (Dylus et al., 2022) using RaxML version 8.1.15 (PROTFAMMAUTO model). Our molecular phylogeny, which matches published species topologies (Rojas et al., 2016; Potter et al., 2021), includes a broad taxonomic sampling, incorporating Miniopteridae, Vespertilionidae, and Molossidae (Figure 2, Branch A; Miller-Butterworth et al., 2007; Lim, 2009). The resulting phylogeny was used for all exploratory selection tests of candidate genes along all branches (Table S3). The tested DNA sequences were from single-copy orthologs found using OMA (Table S4). For selection tests, we used the codon-based method measuring non-synonymous (dN) to synonymous nucleotide (dS) substitutions (the dN/dS metric, ω) with the tool HyPhy (hyphy.org). To identify branches under positive selection, aBSREL (adaptive branch-site random effects likelihood) was used (Smith *et al*. 2015). For the detection of genetic sites under positive selection, MEME (mixed effects model of evolution) was used on all branches (Murrell et al., 2012).

### 3.4 Structural phylogenetics using Alphafold and Foldseek

Alphafold is a large scale language model used to impute protein structures from sequence, which is described in Jumper et al. 2021. We used Alphafold v2 which is available at https://github.com/google-deepmind/alphafold. With the protein structure, we then performed structural alignments with Foldseek, which is described in van Kempen et al. 2023 and can be downloaded at https://github.com/steineggerlab/foldseek. The metric for assessing the topological similarity of protein structure is the TM-score, which has the value in (0,1), where 1 represents a perfect match between two structures.

### 3.5 HCR™ RNA-fluorescent in situ hybridization

Additional glucose tolerance tests (Supp Table 6) were performed on *Pteronotus parnellii* and *Micronycteris minuta* (insect diet), *Carollia perspicillata* (piper-insect diet), *Artibeus jamaicensis* (fig-fruit diet), *Anoura geoffroyi* and *Glossophaga soricina* (nectar diet), and *Phyllostomus discolor* (insect-fruit-nectar diet) in April 2022 and April 2023 with permission from the Republic of Trinidad and Tobago. To determine blood glucose levels, a drop of blood was drawn from the forearm with a 30G lancet before the sugar bolus and 10 minutes post-feeding. The blood was immediately measured using an AlphaTRAK 2.0 glucometer (Zoetis®) with a range of 20-750 mg/dL calibrated for cats (health range 120-300 mg/dL). Finally, the bats were euthanized (n=2-4 per species) and the tissue was dissected, fixed and stored in 4% paraformaldehyde.

Fluorescently labeled HCR hairpins targeting SGLT1 (*Slc5a1*), GLUT2 *(Slc2a2)*, and GLUT5 *(Slc2a5)* in fasted and fed bats were purchased from Molecular Technologies and 3^rd^ generation HCR RNA-FISH was performed on 10 µm paraffin sections of the duodenum. Slides were deparaffinized and microwave antigen retrieval was applied at 95°C for 15 min. A hydrophobic barrier was created using ImmEdge® Hydrophobic Barrier PAP Pen (Vector laboratories) around the sections on a slide, and bleaching solution (3% H2O2, 5% formamide, 0.5X SSC) was applied for 30 min under strong LED light. Tissue sections were permeabilized with Proteinase K (20 ug/mL) for 12 min at room temperature, washed with PBS five times, incubated in hybridization buffer for 30 min at 37°C, then incubated in Hybridization buffer containing probes at 37°C for 16 hours. Tissue sections were washed 5 times with the wash buffer for 5 min each, then 2 times in the 5 X SSCT (5X SSC + 0.1% Tween 20). Fluorescently labeled HCR amplifiers were snap-cooled by heating at 95°C for 90 s and cooling to room temperature for 30 min under dark conditions and added to the amplification buffer. Tissue sections were incubated in the amplification buffer for 30 min at room temperature and incubated in an amplification buffer containing amplifiers overnight at room temperature in a humid chamber under dark conditions. Slides were washed 4 times in 5x SSCT for 5 min each, stained with DAPI (10ug/ml) in 5X SSCT for 30 min, and then washed 2 times in 5X SSCT. Prolong gold (Invitrogen) was mounted, and slides were stored at 4°C until imaging.

### 3.6 Image analysis

Images were taken on a Nikon Ti Eclipse with CSU-W1 spinning disk confocal microscope at 40x magnification. Fiji software was used to manually segment 5 cells per section and each cell was used to quantify fluorescence signal using threshold-based segmentation and spot detection (20-55 per treatment per species). Two sections with good morphology per individual were chosen for study. Data for fluorescence intensity was log2 transformed for comparisons between time points within a species and across species. We performed Games-Howell pairwise comparisons using the *ggstatsplot* function in R.

### 3.7 Quantitative PCR

Total RNA was isolated from 3-5mg flash frozen duodenum tissue from *Ptenonotus parnellii* (n_t=10_=3, n_t=0_=3), *Carollia perspicillata* (n_t=10_=2, n_t=0_=3)*, Artibeus jamaicensis* (n_t=10_=2, n_t=0_=1) *Anoura geoffroyi* (n_t=10_=4, n_t=0_=3) and *Glossophaga soricina* (n_t=10_=3, n_t=0_=1). RNA was extracted using Promega Maxwell RSC simplyRNA Tissue Kit. Quantification was performed using Qubit RNA High Sensitivity Assay Kit and quality was assessed using a 1% agarose gel stained with ethidium bromide. Downstream cDNA synthesis was performed using Applied Biosystems High-Capacity RNA-to-cDNA Kit using a programmed thermal cycler at 37°C for 60 minutes, followed by a 5 minute incubation at 95°C, before cooling to 4°C. The specific primer sequences used for target genes SGLT1 (*Slc5a1*), GLUT2 *(Slc2a2)* Sucrase-Isomaltase (*SI)*, and reference gene GAPDH are shown in Supplementary Table 7A and were ordered from Integrated DNA Technologies (IDT - Carolville, IA). Primer pair validation and experimentation was performed using QuantStudio 7 Pro Real-time qPCR platform and software. Data generated from qPCR was analyzed using the 2^-ΔΔCt^ method (Livak & Schmittgen, 2001) to assess variance in relative fold gene expression between SGLT1, SI, and GLUT2 against the housekeeping gene GAPDH. For heatmap visualization of normalized gene expression (ΔCt), the R package ‘pheatmap’ version 1.0.12 was used. Clusters are based on ‘euclidean’ distance measures, also performed on the R package ‘pheatmap’. Statistical analysis on fold gene expression changes (ΔΔCt) at t=10 relative to t=0 between species was evaluated with a one-way ANOVA in GraphPad Prism 10.2.0 (Supplementary Table 7B).

### 4.1 Data depository

Original data underlying this manuscript can be accessed from the Stowers Original Data Repository at http://www.stowers.org/research/publications/LIBPB-2406. Original genome data and analysis can be found here: https://simrbase.stowers.org/jb_pub/

## Acknowledgements

A portion of the fieldwork was carried out within the framework of the project “Contribución a la conservación del Bosque seco Tropical del Valle del Cauca’’ under the CVC permit No. 1122 of 2018, from the Institute for Research and Preservation of the Cultural and Natural Heritage of Valle del Cauca (INCIVA), whom we thank, along with the support of Jose Omar Ortíz, Ruth Rivera and Andrés Bernal, and the field assistance from Therios (Cali, Colombia), Cristian Calvache, Sergio Tabares, Alejandro Chito and Oscar Cuellar. Thanks to Maria Eifler at the KU Biodiversity Institute and Natural History Museum, Nancy Simmons, and Marisa Surovy from the AMNH for facilitating access and research of museum specimens. We thank the scientists at the Stowers Institute for Medical Research scientists, Seth Malloy, Hannah Wilson, Michael Frangello, Cindy Maddera, and Xia Zhao, for help in processing and imaging histological and EM sections as well as Dan Bradford for providing training on RT-qPCR instrumentation and data analysis. We also thank Jose Alejandro Riascos for the bat illustration in the graphical abstract, Darshan Narang, Richard Smith, and Shivam Mahadeo for their support in Trinidad, and Fiona Reid and Jerrica Jamison for her support in Costa Rica, as well as Sharlene Sanatana for guidance in PCM analysis, Alexa Sadier and Laurel Yohe for comments that improved this manuscript. Finally, we thank the Rohner lab members for creating a stimulating scientific environment and for their support during the manuscript writing. Funding for the project was provided by the SIMR to N.R., the NSF Postdoctoral Research Fellowships in Biology #2109717 (2021-2022) to J.C., the BWF Postdoctoral Diversity Enrichment Program (G-1022339) to J.C, and the HHMI Hanna H. Gray Fellows Program (GT15991) to J.C. Any opinions, findings, and conclusions or recommendations expressed in this material are those of the authors and do not necessarily reflect the views of the NSF, BWF, or HHMI.

## Author contributions

A.B.R. and J.C. designed the study. A.B.R., J.C., and V.P. designed the figures. A.B.R. performed the *in vivo* physiology experiments as an undergraduate thesis project (2019-2020) directed by O.E.M.G. A.B.R. and O.E.M.G. designed the *in vivo* physiology experiments and performed the statistical analyses. A.B.R. and J.C. performed additional glucose tolerance tests and obtained gut tissue for molecular analyses in April 2022. A.B.R. and J.C. dissected the gastrointestinal tract of museum bats and performed morphological data interpretation. K.Y. performed the TEM experiments and imaging. A.B.R. made the gut anatomy measurements and comparisons. S.R. produced gene models across the assembled genomes, assigned orthologs, built webpages, and genome browsers, and generated the phylogenomic tree. J.R. and J.C. performed selection tests. J.R. performed Alphafold and Folkseek analysis. J.C., D.T., and Y.W. performed the HCR experiments, J.C. and V.P. performed HCR imaging, and J.C. performed HCR analysis. V.P. and P.M. extracted RNA from the duodenum and P.M. performed qPCR. A.B.R., J.C., and N.R. co-wrote the manuscript.

## Notes

https://simrbase.stowers.org/bats/data

### Competing Interest Statement

The authors have declared no competing interest.

### Summary of Updates

More bat species included; new results; Figure 2 revised; Figure 5 revised; added a new Figure 6

