## Extended Data for "Sugar assimilation underlying dietary evolution of Neotropical bats"

#### Supplemental information/Extended Data

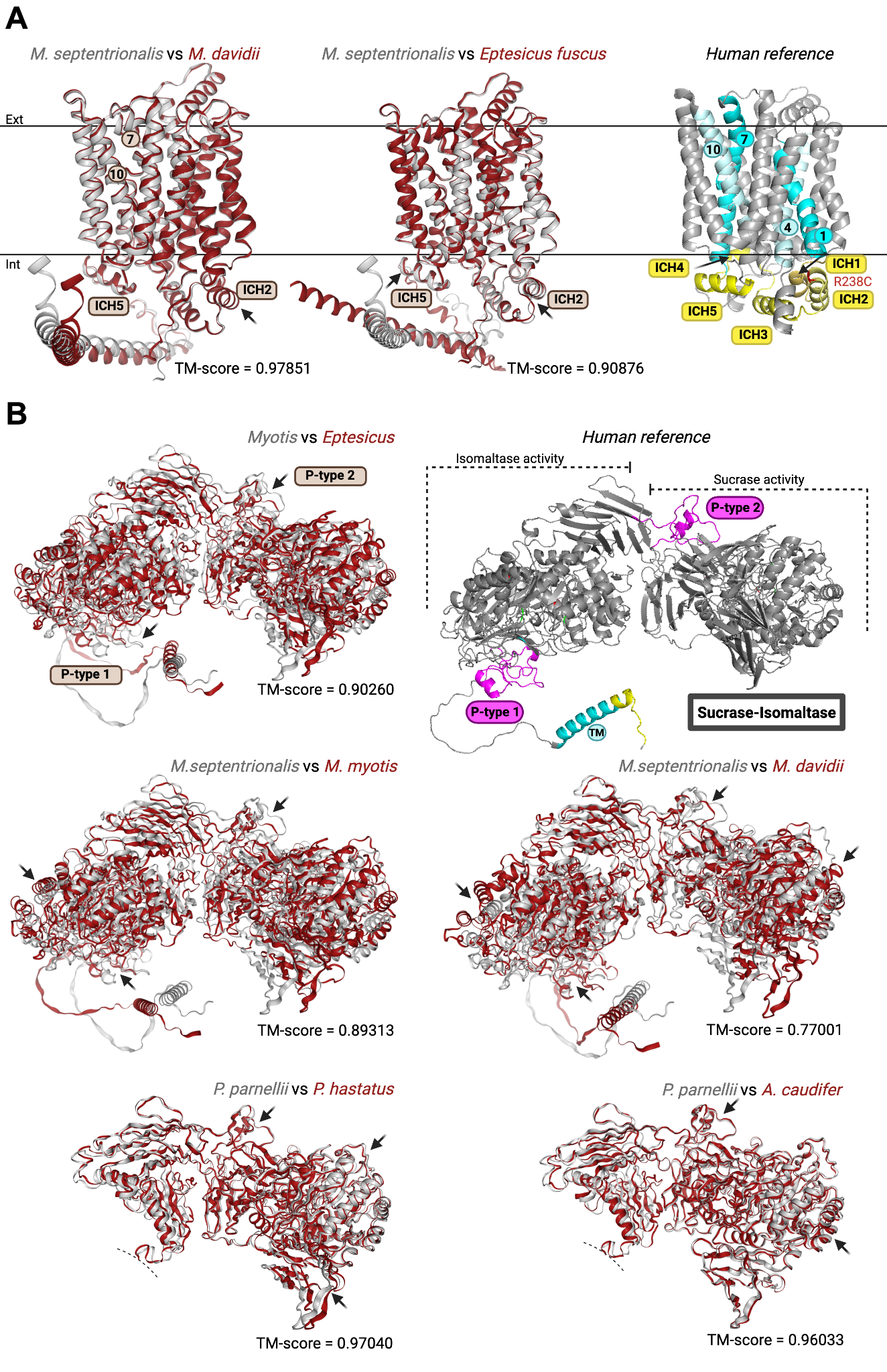

**Supplemental Figure 1.** Structural changes of proteins encoded by genes under positive selection in the focal branch. A) SLC2A5 within *Myotis* species and between *Myotis* to their sister genus *Eptesicus*. The intracellular regions highlighted with an arrow focuses on the observed structural changes, summarized as a TM-score. TM-score = 1 is a perfect structural match. B) Folkseek comparisons for sucrase-isomaltase (SI) of *Myotis*, *Phyllostomus*, and *Anoura.* The observed variation in SI from *Myotis* species indicates there may be functional changes to both the isomaltase and sucrase enzyme properties. The human reference protein structure has additional features highlighted for protein orientation as follow: transmembrane (TM) 1 and 4 in the N-terminal domain are colored in cyan and light cyan, respectively; TM 7 and 10 in the C-terminal bundle are colored in cyan and light cyan, respectively; intracellular domain helices (ICH) unique to the sugar transporters are shown in yellow; single amino acid change (R238C) related to colon cancer. For SI, P-type 1 and 2 are colored in magenta, the binding sites for sugar are colored in green, and the known mutations that affect function are colored in red. Protein annotations were made using PyMOL.

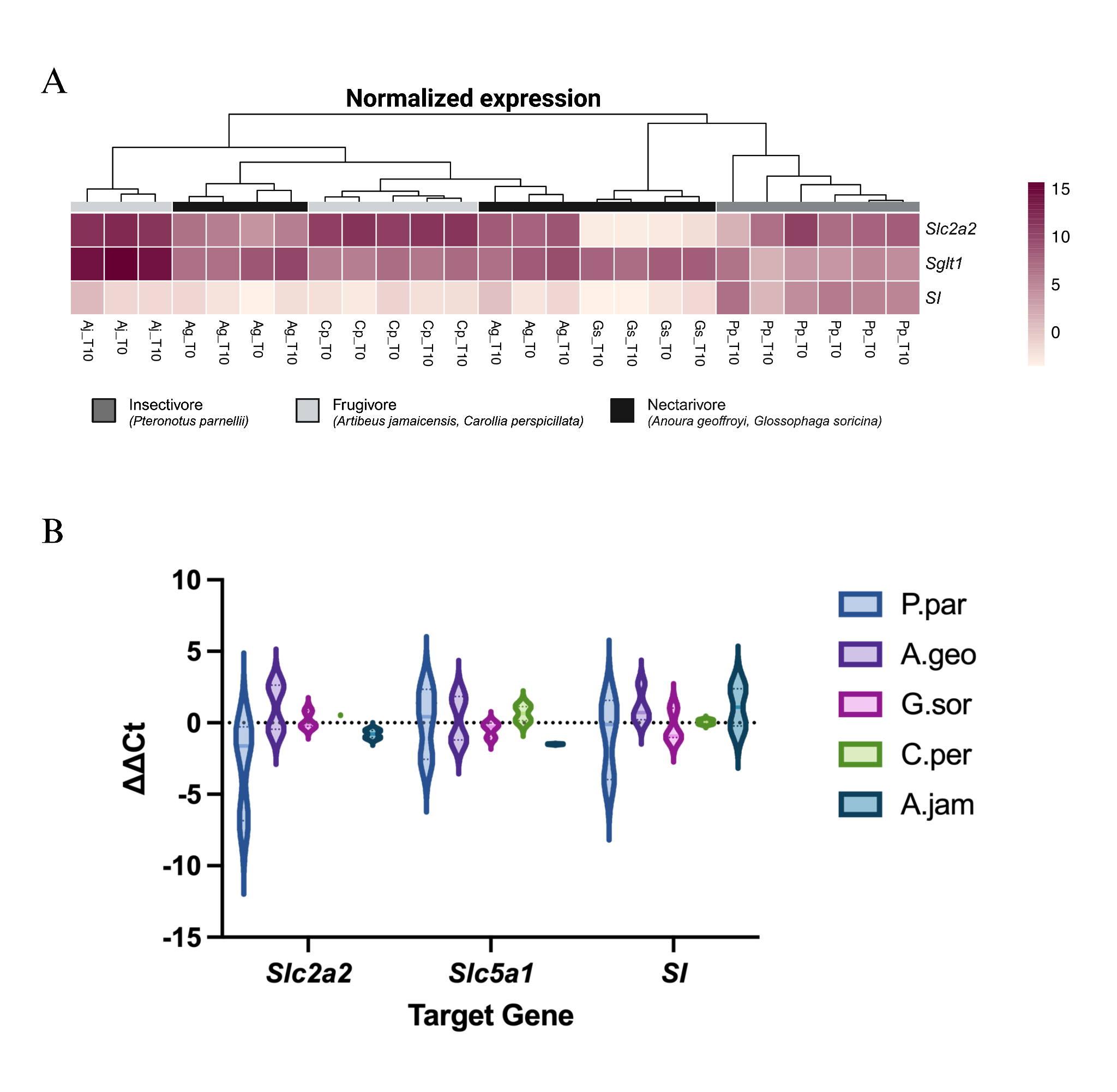

**Supplemental Figure 2.** (A) RT-qPCR heatmap of normalized expression (ΔCt) relative to housekeeping gene GAPDH of nutrient transporters *Slc2a2* (GLUT2), *Slc5a1* (SGLT1) and digestive enzyme *SI* (SI). Five species representing four dietary guilds are shown [insects, *Ptenonotus parnellii* (Pp, n_t=10_=3, n_t=0_=3); omnivore, *Carollia perspicillata* (Cp, n_t=10_=2, n_t=0_=3); fruit, *Artibeus* *jamaicensis* (Aj, n_t=10_=2, n_t=0_=1); nectar, *Anoura* *geoffroyi* (Ag, n_t=10_=4, n_t=0_=3) and *Glossophaga* *soricina* (Gs, n_t=10_=3, n_t=0_=1)]. Samples are ordered by hierarchical clustering using Euclidean distance and the colors assigned to the clusters correspond to the dietary guild. (B) RT-qPCR-based expression analysis of *Slc2a2*, *Slc5a1* and *SI* fold change at t=10 relative to t=0 (ΔΔCt). Significant differences in fold change expression between species were determined by one-way ANOVA (see Supplementary Table 8B for statistical details).
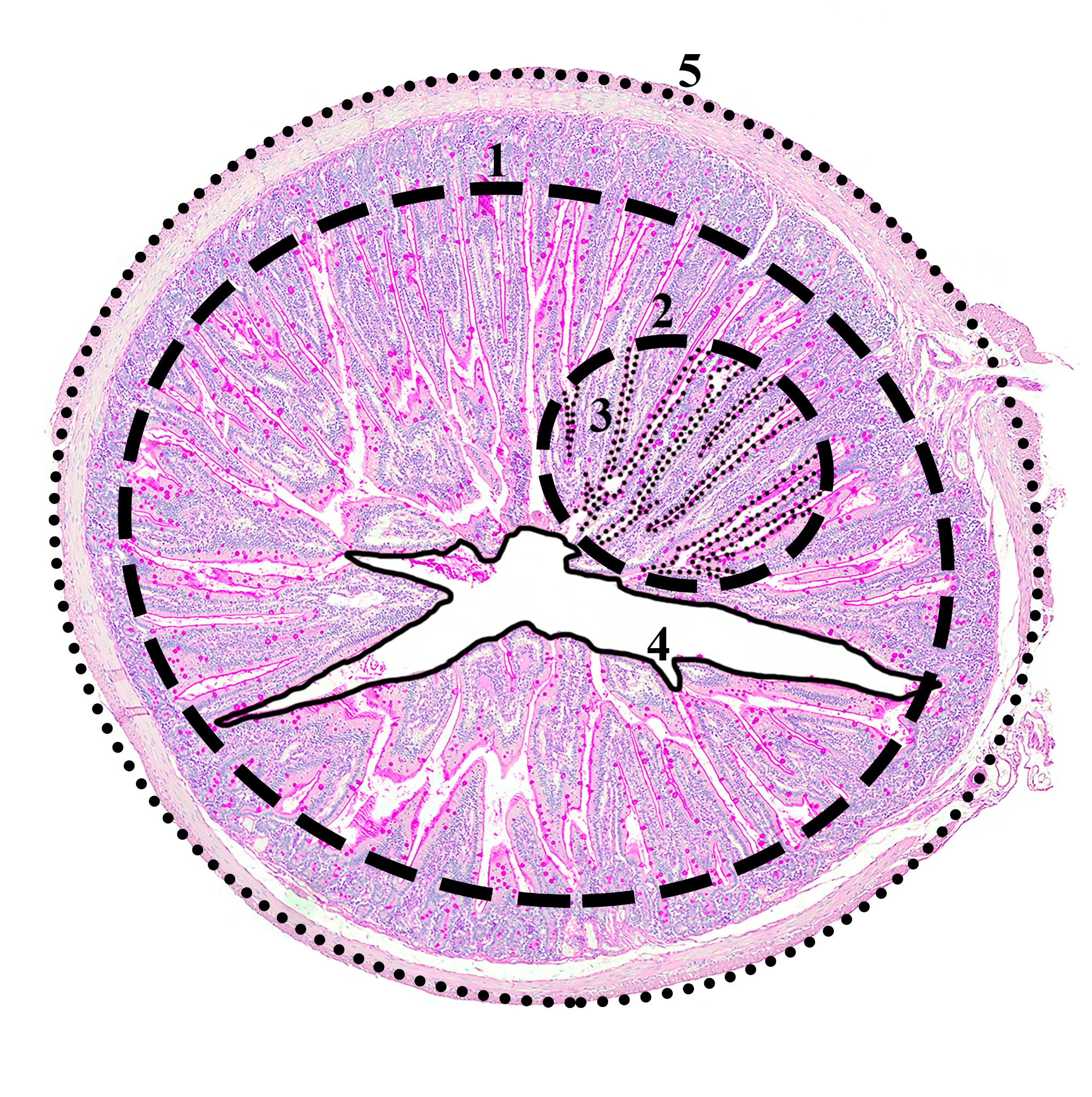

**Supplemental Figure 3.** Measures extracted from the intestine cross sections. 1. Villi-lumen area (VLA); 2. Sample area (SA); 3. Villi perimeter in the sampled area (VPSA); 4. Lumen area (LA); 5. Cross section perimeter (CSP).

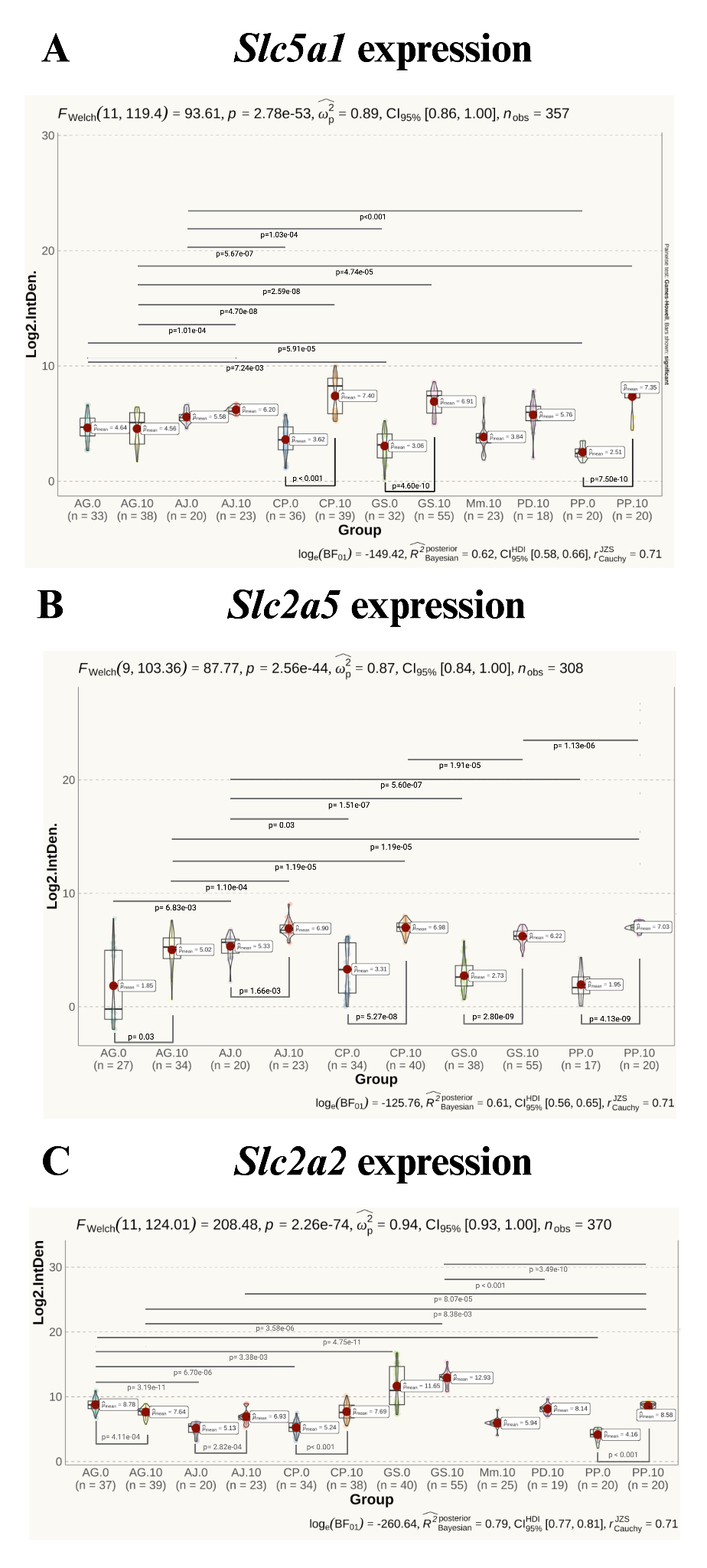

**Supplemental Figure 4.** HCR RNA-FISH data for *Slc5a1* (A), *Slc2a5* (B), and *Slc2a2* (C). Comparisons within species at t=0 and t=10 are at the bottom of each graph. Comparisons between species at each respective time point are shown at the top of each graph. The following species represent each dietary guild: insects, *Ptenonotus parnellii* (PP, n=4) and *Micronycteris minuta* (Mm n=1); omnivore, *Carollia perspicillata* (CP, n=4); fruit, *Artibeus* *jamaicensis* (AJ, n=2); nectar, *Glossophaga* *soricina* (GS, n=4) and *Anoura* *geoffroyi* (AG, n=4). *Phyllostomus discolor* (PD, n=1), is an omnivorous bat that has a large portion of nectar in their diet. Only significant data (p_adj_ < 0.001) are shown. The number of enterocytes analyzed are shown below each group.

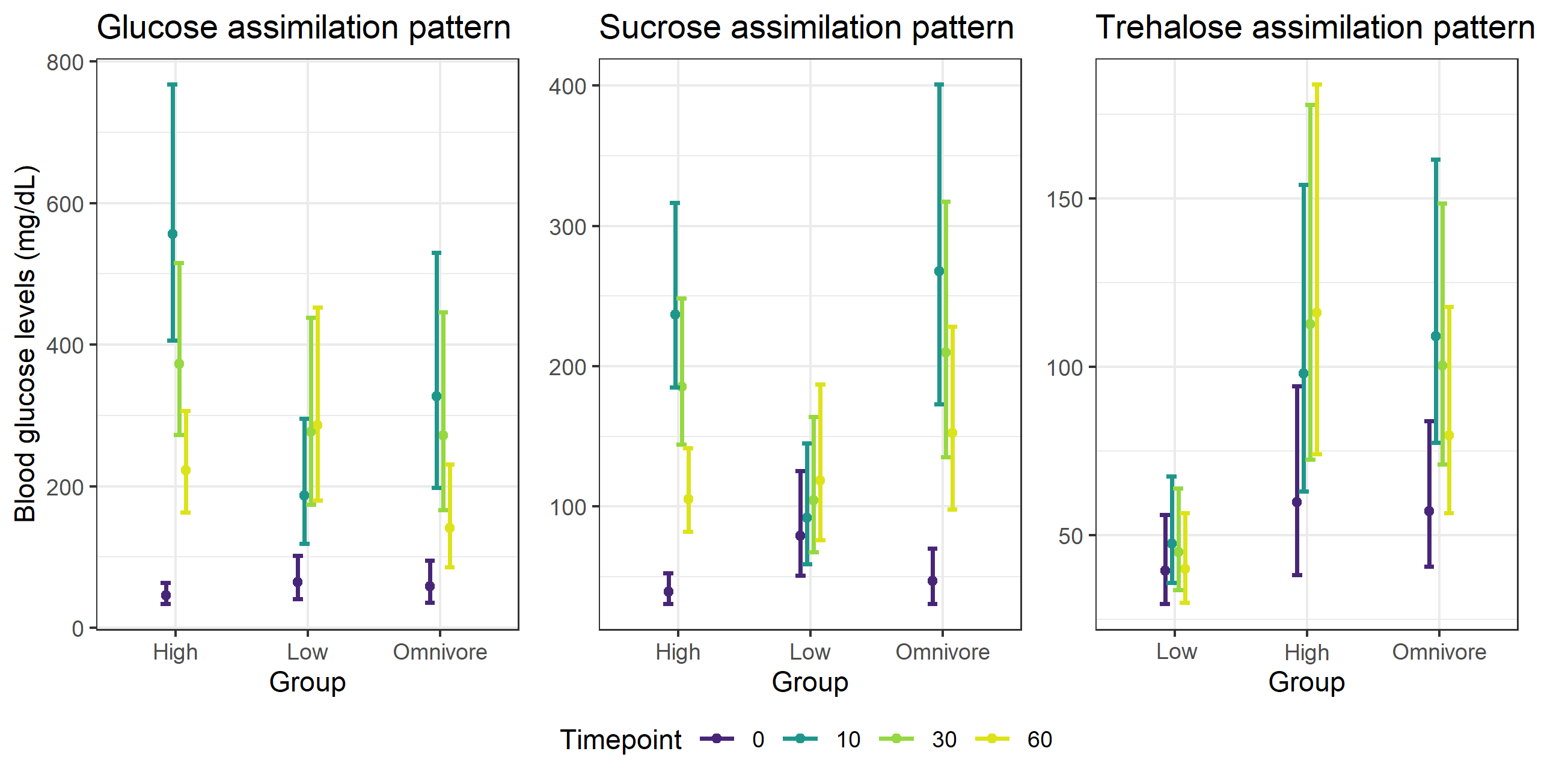

**Supplemental Figure 5.** Comparison of assimilation curves for glucose, sucrose, and trehalose solutions among Neotropical bats with different food preferences: high sugar in glucose and sucrose graphs refer to frugivorous and nectarivorous bats, while low sugar refers to insectivorous, carnivorous and hematophagous bats; high sugar in the trehalose graph refers to insectivorous bats and low sugar refers to the rest of the dietary categories; omnivores have their own category because they have diets where the three sugars are present in relatively high proportions.

#### Supplemental Tables

**Supplemental Table 1.** Species included in the *in vivo* physiology essay. *A. lit: Artibeus lituratus, A. pla: Artibeus planirostris, A. aeq: Artibeus aequatorialis, D. pha: Dermanura phaeotis, U. con: Uroderma convexum, U. bak: Uroderma bakeri, P. hel: Platyrrhinus helleri, S. gia: Sturnira giannae, S. lud: Sturnira ludovici, S. lui: Sturnira luisi, S. par: Sturnira parvidens, G. sp.: Glossophaga* sp.*, C. god: Choeroniscu godmani, L. con: Lonchophylla concava, L. rob: Lonchophylla robusta, C. cas: carollia castanea, C. per: Carollia perspicillata, C. bre: Carollia brevicauda, P. dis: Phyllostomus discolor, P. has: Phyllostomus hastatus, V. spe: Vampyrum spectrum, D. rot: Desmodus rotundus, G. cre: Gardnerycteris crenulatum, M. cau: Myotis caucensis, M. alb: Myotis albescens, M. mol: Molossus molossus, S. bil: Saccopteryx billineata, P. kap: Peropteryx kappleri.*

| **Family** | **Genus** | **Food preference** | **Sugar** | **# Individuals** | **Species** |
| --- | --- | --- | --- | --- | --- |
| Phyllostomidae | *Artibeus* | Fruits | Trehalose | 17 | *A. lit, A. pla, A. aeq* |
|  |  |  | Sucrose | 22 |  |
|  |  |  | Glucose | 13 |  |
|  | *Dermanura* |  | Trehalose | 6 | *D. pha* |
|  |  |  | Sucrose | 6 |  |
|  |  |  | Glucose | 6 |  |
|  | *Uroderma* |  | Trehalose | 1 | *U. con, U. bak* |
|  |  |  | Sucrose | 1 |  |
|  |  |  | Glucose | 1 |  |
|  | *Platyrrhinus* |  | Sucrose | 1 | *P. hel* |
|  | *Sturnira* |  | Trehalose | 1 | *S. gia, S. lud, S. lui, S. par* |
|  |  |  | Sucrose | 1 |  |
|  |  |  | Glucose | 3 |  |
|  | *Glossophaga* | Nectar | Trehalose | 4 | *G. sp.* |
|  |  |  | Sucrose | 6 |  |
|  |  |  | Glucose | 4 |  |
|  | *Choeroniscus* |  | Trehalose | 1 | *C. god* |
|  |  |  | Sucrose | 1 |  |
|  |  |  | Glucose | 2 |  |
|  | *Lonchophylla* |  | Glucose | 5 | *L. con, L. rob* |
|  | *Carollia* | Fruits & Insects | Trehalose | 18 | *C. cas, C. per, C. bre* |
|  |  |  | Sucrose | 19 |  |
|  |  |  | Glucose | 13 |  |
|  | *Phyllostomus* | Insects, Nectar, Polen, Fruits, Vertebrates | Trehalose | 3 | *P. dis, P. has* |
|  |  |  | Sucrose | 1 |  |
|  |  |  | Glucose | 1 |  |
|  | *Vampyrum* | Vertebrates | Glucose | 1 | *V. spe* |
|  | *Desmodus* | Blood | Trehalose | 4 | *D.rot* |
|  |  |  | Sucrose | 4 |  |
|  |  |  | Glucose | 6 |  |
|  | *Gardnerycteris* | Insects | Trehalose | 1 | *G. cre* |
| Mormoopidae | *Pteronotus* |  | Glucose | 12 | *P. par* |
| Vespertilionidae | *Myotis* |  | Trehalose | 3 | *M. cau, M. alb* |
|  |  |  | Sucrose | 1 |  |
|  |  |  | Glucose | 2 |  |
| Molossidae | *Molossus* |  | Trehalose | 1 | *M. mol* |
|  |  |  | Glucose | 1 |  |
| Emballonuridae | *Saccopteryx* |  | Trehalose | 1 | *S. bil* |
|  |  |  | Sucrose | 1 |  |
|  |  |  | Glucose | 1 |  |
|  | *Peropteryx* |  | Sucrose | 3 | *P. kap* |
| **Total** | | | | **199** | **29** |

**Supplemental** **Table 2.** Phylogenetic signal evaluated for the assimilation proxy (corrected area under the curve) of glucose, sucrose and trehalose in Neotropical bats.

| **Phylogenetic signal** | | | |
| --- | --- | --- | --- |
| **Sugar** | **Variable** | **Lambda (λ)** | **P-value** |
| Glucose | Assimilation proxy | 0.988169 | 0.0584431 |
| Sucrose | Assimilation proxy | **0.999927** | **0.0158425** |
| Trehalose | Assimilation proxy | 0.70459 | 0.281583 |

**Supplemental Table 3**. Twenty-two bat genomes were assembled and annotated for selection tests. Species were selected to best match the diversity of the *in vivo* physiology data. Two outgroup genomes, human and shrew, were used to polarize the evolutionary changes in bats. In addition to the twenty-two Neotropical bat genomes, we obtained an additional 24 genomes from all available bat species for structural comparisons of proteins with Foldseek.

| **Family** | **Species** | **Year published** | **NCBI accession** | **RNAseq/ Proteome** | **Species Code** |
| --- | --- | --- | --- | --- | --- |
| Vespertilionidae | *Eptesicus fuscus* | 2020 | *DNAzoo SRS7189612 | GCF_000308155.1 | EFU |
| Vespertilionidae | *Myotis myotis* | 2020 | GCA_014108235.1 | GCF_014108235.1 | MMY |
| Vespertilionidae | *Myotis septentrionalis* | 2020 | *DNAzoo  SRS7189622 | Hi-C | MSE |
| Molossidae | *Molossus molossus* | 2020 | GCA_014108415.1 | GCF_014108415.1, SRR9703456 | MMO |
| Miniopteridae | *Miniopterus natalensis* | 2016 | GCA_001595765.1 | GCF_001595765.1 | MNA |
| Miniopteridae | *Miniopterus schreibersii* | 2019 | GCA_004026525.1 | n/a | MSC |
| Mormoopidae | *Pteronotus parnellii* | 2013 | GCA_000465405.1 | SRR9703452, PRJNA481095, PRJNA481095 | PPA |
| Mormoopidae | *Mormoops blainvillei* | 2019 | GCA_004026545.1 | SRR9703451 | MBL |
| Phyllostomidae | *Desmodus rotundus* | 2018 | GCA_002940915.1 | GCF_002940915.1, SRR9703482, PRJNA178123, SRR8878915 | DRO |
| Phyllostomidae | *Anoura caudifer* | 2019 | GCA_004027475.1 | SRR9703485 | ACA |
| Phyllostomidae | *Musonycteris harrisonii* | 2021 | *Gigadb 100746 | n/a | MHA |
| Phyllostomidae | *Leptonycteris nivalis* | 2020 | [PRJNA543656](https://www.ncbi.nlm.nih.gov/sra?linkname=bioproject_sra_all&from_uid=543656) | n/a | LNI |
| Phyllostomidae | *Leptonycteris yerbabuenae* | 2020 | PRJNA627035 | SRR9087861, SRR9087862, SRR9087863, SRR9087864 | LYE |
| Phyllostomidae | *Macrotus waterhousii* | 2020 | *Gigadb 100746 | SRR9703483 | MWA |
| Phyllostomidae | *Macrotus californicus* | 2019 | GCA_007922815.1 | n/a | MCA |
| Phyllostomidae | *Micronycteris hirsuta* | 2019 | GCA_004026765.1 | n/a | MHI |
| Phyllostomidae | *Tonatia saurophila* | 2019 | GCA_004024845.1 | n/a | TSA |
| Phyllostomidae | *Phyllostomus discolor* | 2019 | GCA_004126475.2 | GCF_004126475.2 | PDI |
| Phyllostomidae | *Phyllostomus hastatus* | 2021 | GCA_019186645.1 | SRR9703480 | PHA |
| Phyllostomidae | *Sturnira hondurensis* | 2020 | GCA_014824575.2 | GCF_014824575.2 | SHO |
| Phyllostomidae | *Carollia perspicillata* | 2019 | GCA_004027735.1 | SRR9703464, SRR5872538 | CPE |
| Phyllostomidae | *Artibeus jamaicensis* | 2019 | GCA_004027435.1 | SRR9703479, SRR539297, PRJNA305413 | AJA |
| Primates | *Homo sapiens* | 2019 | GCA_000001405.28 | GCF_000001405.39 | HSA |
| Eulipotyphla | *Sorex araneus* | 2012 | GCA_000181275.2 | n/a | SAR |

| **Family** | **Species** | **Year published** | **NCBI accession** | **Species Code** |
| --- | --- | --- | --- | --- |
| Vespertilionidae | *Myotis brandtii* | 2013 | GCF_000412655.1 | MYB |
| Vespertilionidae | *Myotis davidii* | 2012 | GCF_000327345.1 | MDA |
| Vespertilionidae | *Murina aurata feae* | 2019 | GCA_004026665.1 | MAU |
| Molossidae | *Tadarida brasiliensis* | 2019 | GCA_004025005.1 | TBR |
| Vespertilionidae | *Aeorestes cinereus* | 2020 | GCA_011751065.1 | ACI |
| Vespertilionidae | *Lasiurus borealis* | 2019 | GCA_004026805.1 | LBO |
| Vespertilionidae | *Pipistrellus kuhlii* | 2020 | GCF_014108245.1 | PKU |
| Vespertilionidae | *Pipistrellus pipistrellus* | 2020 | GCA_903992545.1 | PPI |
| Pteropodidae | *Pteropus alecto* | 2013 | GCF_000325575.1 | PAL |
| Pteropodidae | *Pteropus rufus* | 2020 | SRR11097142 | PRU |
| Pteropodidae | *Pteropus alecto* | 2013 | GCF_000325575.1 | PGI |
| Pteropodidae | *Rousettus aegyptiacus* | 2020 | GCF_014176215.1 | RAE |
| Pteropodidae | *Rousettus madagascariensis* | 2019 | SRR11097137 | RMA |
| Pteropodidae | *Eonycteris spelaea* | 2018 | GCA_003508835.1 | ESP |
| Pteropodidae | *Macroglossus sobrinus* | 2019 | GCA_004027375.1 | MSO |
| Pteropodidae | *Eidolon helvum* | 2013 | GCA_000465285.1 | EHE |
| Pteropodidae | *Eidolon dupreanum* | 2020 | ASM46528v1_HiC | EDU |
| Hipposideridae | *Hipposideros armiger* | 2016 | GCF_001890085.1 | HAR |
| Hipposideridae | *Hipposideros galeritus* | 2019 | GCA_004027415.1 | HGA |
| Rhinolophidae | *Rhinolophus ferrumequinum* | 2019 | GCA_007922735.1 | RFE |
| Megadermatidae | *Megaderma lyra* | 2019 | GCA_004026885.1 | MLY |
| Rhinolophidae | *Craseonycteris thonglongyai* | 2019 | GCA_004027555.1 | CTH |
| Noctilionidae | *Noctilio leporinus* | 2019 | GCA_004026585.1 | NLE |

**Supplemental Table 4.** Statistical summary of the exploratory aBSREL results. aBSREL (adaptive Branch-Site Random Effects Likelihood) will test whether a proportion of sites have evolved under positive selection along each branch in the phylogeny. Baseline model refers to MG94xREV baseline model that infers a single omega rate per branch. Full adaptive model infers an optimized number of omega rate categories per branch. After aBSREL fits the full adaptive model, the Likelihood Ratio Test (LRT) is performed at each branch and compares the full model to a null model where branches are not allowed to have rate classes of ω>1.

| **Gene** | **aBSREL Model** | **AIC_C_** | **logL** | **Parameters** |
| --- | --- | --- | --- | --- |
| *Slc2a1* | Nucleotide GTR | 32127.80 | -16014.83 | 49 |
|  | Baseline | 30189.65 | -15002.06 | 92 |
|  | Full adaptive model | 29812.02 | -14805.10 | 100 |
| *Slc2a2* | Nucleotide GTR | 74075.49 | -36984.67 | 53 |
|  | Baseline | 72399.71 | -36117.31 | 82 |
|  | Full adaptive model | 72101.26 | -35951.86 | 98 |
| *Slc2a3* | Nucleotide GTR | 44911.77 | -22402.81 | 53 |
|  | Baseline | 43351.46 | -21574.92 | 100 |
|  | Full adaptive model | 42565.78 | -21153.57 | 128 |
| *Slc2a4* | Nucleotide GTR | 49387.03 | -24640.44 | 53 |
|  | Baseline | 47689.57 | -23744.00 | 100 |
|  | Full adaptive model | 47653.74 | -23724.05 | 102 |
| *Slc2a5* | Nucleotide GTR | 49623.49 | -24760.68 | 51 |
|  | Baseline | 46931.60 | -23379.26 | 86 |
|  | Full adaptive model | 46529.65 | -23158.00 | 106 |
| *Slc5a1* | Nucleotide GTR | 93090.21 | -46494.05 | 51 |
|  | Baseline | 89484.54 | -44653.76 | 88 |
|  | Full adaptive model | 88528.46 | -44155.46 | 108 |
| *SI* | Nucleotide GTR | 195876.34 | -97887.15 | 51 |
|  | Baseline | 191316.64 | -95568.12 | 90 |
|  | Full adaptive model | 189682.39 | -94710.79 | 130 |
| *Treh* | Nucleotide GTR | 14800.36 | -7375.14 | 25 |
|  | Baseline | 14468.32 | -7185.76 | 48 |
|  | Full adaptive model | 14453.92 | -7174.49 | 52 |

**Supplemental Table 5.** Detailed aBSREL significant results in bats. The results are from exploratory analysis where all branches are tested for positive selection. In this scenario, p-values at each branch must be corrected for multiple testing (using the Holm-Bonferroni correction). The Likelihood Ratio Test (LRT) is performed at each branch and compares the full model to a null model where branches are not allowed to have rate classes of ω>1. Branches and nodes under selection are shown below and correspond to **Figure 2.**

| **Gene** | **Branch** | **LRT** | **p-value** | **𝞈 distribution** |
| --- | --- | --- | --- | --- |
| *Slc2a1* | ACA | 170.2147 | <0.0001 | ω_1_= 0.00 (35%)  ω_2_= 9090 (65%) |
|  | Node 30 | 13.0492 | 0.0005 | ω_1_= 0.00 (99%)  ω_2_= 111 (0.54%) |
| *Slc2a2* | ACA | 30.9776 | <0.0001 | ω_1_= 1.00 (37%)  ω_2_= 334 (63%) |
|  | MHA | 64.2921 | <0.0001 | ω_1_= 1.00 (20%)  ω_2_= 1e5 (80%) |
|  | Node 13 | 35.4885 | <0.0001 | ω_1_= 0.00 (33%)  ω_2_= 1e5 (67%) |
|  | MMY | 19.7098 | <0.0001 | ω_1_= 0.794 (12%)  ω_2_= 80.6 (88%) |
| *Slc2a3* | DRO | 30.6385 | <0.0001 | ω_1_= 0.104 (98%)  ω_2_= 247 (1.9%) |
|  | MSE | 302.4789 | <0.0001 | ω_1_= 0.00 (46%)  ω_2_= 868 (54%) |
|  | Node 10 | 39.5755 | <0.0001 | ω_1_= 0.272 (98%)  ω_2_= 1e5 (1.7%) |
|  | Node 36 | 29.7309 | <0.0001 | ω_1_= 0.00 (90%)  ω_2_=11.6 (9.6%) |
|  | SAR | 15.3250 | 0.0002 | ω_1_= 0. (72%)  ω_2_= 0.611 (14%)  ω_3_= 1e5 (13%) |
|  | Node 37 | 13.3182 | 0.0004 | ω_1_= 0.374 (99%)  ω_2_= 158 (0.67%) |
| *Slc2a4* | n/a | n/a | n/a | n/a |
| *Slc2a5* | EFU | 28.7577 | <0.0001 | ω_1_= 0.243 (11%)  ω_2_= 13.9 (89%) |
|  | MSE | 15.5125 | 0.0001 | ω_1_= 0.351 (93%)  ω_2_= 1e5 (6.7%) |
|  | Node 39 | 12.1544 | 0.0008 | ω_1_= 0.024 (3.7%)  ω_2_= 13.2 (96%) |
| *Slc5a1* | MNA | 447.4476 | <0.0001 | ω_1_= 0.0776 (90%)  ω_2_=100000 (10%) |
|  | TSA | 16.1794 | 0.0001 | ω_1_= 1e9 (100%) |
| *SI* | MNA | 96.3819 | <0.0001 | ω_1_= 1.00 (98%)  ω_2_= 1.00 (1.8%)  ω_3_= 1e5 (0.60%) |
|  | MSC | 53.7631 | <0.0001 | ω_1_= 0.301 (99%)  ω_2_= 6250 (0.5%) |
|  | MWA | 27.5404 | <0.0001 | ω_1_= 0.00 (2.9%)  ω_2_= 1e5 (97%) |
|  | Node 34 | 45.6658 | <0.0001 | ω_1_= 1.00 (6.3%)  ω_2_= 9090 (94%) |
|  | Node 39 | 38.1848 | <0.0001 | ω_1_= 0.215 (97%)  ω_2_= 40.7 (3.1%) |
|  | PHA | 49.2379 | <0.0001 | ω_1_= 1.00 (3.9%)  ω_2_= 3850 (96%) |
|  | MSE | 20.0862 | <0.0001 | ω_1_= 0.200 (5%)  ω_2_= 1e5 (95%) |
|  | Node 10 | 17.5613 | 0.0001 | ω_1_= 0.00 (92%)  ω_2_= 7.81 (8%) |
| *Treh* | Node 11 | 22.4132 | <0.0001 | ω_1_= 3.77 (100%) |
|  | EFU | 0.0221 | 0.0014 | ω_1_= 0.00 (78%)  ω_2_= 13.2 (22%) |

**Supplemental Table 6.** Change in gene expression and blood glucose levels 10-minutes after eating a 20% glucose solution. Each bat was fed a single dose of glucose (5.4mg/kg body weight) after fasting. Samples in bold were used for HCR FISH, while others were sampled non-lethally.

| Species | ID | Blood glucose (mg/dL)  (t=0) | Blood glucose (mg/dL)  (t=10) | AVG  Blood glucose change ± SE  (mg/dL) | AVG  Log2 fold change  *Slc5a1* | AVG  Log2 fold change  *Slc2a2* | AVG  Log2 fold change *Slc2a5* |
| --- | --- | --- | --- | --- | --- | --- | --- |
| *Carollia perspicillata* | **TT22- 57** | 75 | NA | 440.6 ± 66.72 | 1.102 | 0.468 | 1.11 |
|  | **TT22-45** | 60 | 538 |  |  |  |  |
|  | **TT22-60** | 70 | NA |  |  |  |  |
|  | **TT22-59** | 83 | 343 |  |  |  |  |
|  | TT22-83 | 54 | 719 |  | | | |
|  | TT22-01 | 78 | 342 |  |  |  |  |
|  | TT22-03 | 70 | 606 |  |  |  |  |
| *Glossophaga soricina* | **TT22- 47** | 91 | NA | 659.2 ± 29.27 | 1.26 | 0.109 | 1.277 |
|  | **TT22-51** | 28 | NA |  |  |  |  |
|  | **TT22-50** | 63 | 750 |  |  |  |  |
|  | **TT22-52** | 59 | 707 |  |  |  |  |
|  | TT22-53 | 29 | 750 |  | | | |
|  | TT22-54 | 67 | 597 |  |  |  |  |
|  | TT22-62 | 40 | 750 |  |  |  |  |
| *Pteronotus parnellii* | **TT22-19** | 57 | NA | 148.2 ± 48.82 | 1.813 | 1.062 | 2.605 |
|  | **TT22-28** | 41 | NA |  |  |  |  |
|  | **TT22-21** | 95 | 211 |  |  |  |  |
|  | **TT22-27** | 61 | 293 |  |  |  |  |
|  | TT22-20 | 59 | 241 |  | | | |
|  | TT22-26 | 46 | 136 |  |  |  |  |
|  | TT22-29 | 64 | 185 |  |  |  |  |
| *Anoura*  *geoffroyi* | **TT22-38** | 61 | 750 | 639.4 ± 20.96 | 0.331 | -0.123 | 1.719 |
|  | **TT22-31** | 75 | 742 |  |  |  |  |
|  | **TT22-37** | 70 | NA |  |  |  |  |
|  | **TT22-30** | 66 | NA |  |  |  |  |
|  | TT22-40 | 68 | 750 |  | | | |
|  | TT22-34 | 65 | 638 |  |  |  |  |
|  | TT22-33 | 51 | 586 |  |  |  |  |
| *Phyllostomus disculor* | **TT23-34** | 102 | 750 | 500.2 ± 107.54 | NA | NA | NA |
|  | TT22-64 | 121 | 750 |  | | | |
|  | TT23-55 | 52 | 497 |  |  |  |  |
|  | TT23-46 | 73 | 750 |  |  |  |  |
|  | TT22-101 | 96 | 198 |  |  |  |  |
| *Artibeus jamaicensis* | **TT22-68** | 69 | 243 | 483 ± 96.38 | 0.006 | 0.350 | 0.294 |
|  | **TT22-63** | 37 | NA |  |  |  |  |
|  | TT22-81 | 91 | 750 |  | | | |
|  | TT22-86 | 29 | 632 |  |  |  |  |
|  | TT22-80 | 41 | 328 |  |  |  |  |
|  | TT22-66 | 58 | 750 |  |  |  |  |
| Micronycteris minuta | **TT22-15** | 60 | 412 | 174.7 ± 94.57 | NA | NA | NA |
|  | TT22-72 | 83 | 112 |  | | | |
|  | TT22-48 | 149 | 292 |  |  |  |  |

**Supplementary Table 7:** (A) RT-qPCR samples ran in triplicate against each target gene. Resulting Ct values from QuantStudio 7 Pro Real-time qPCR platform were averaged and ΔCt values were generated from the change between each target gene compared to GAPDH and visualized using heatmap (Supplemental Figure 2A). (B) Within each species tested, t=10 ΔCt values were compared against each t=0 ΔCt value. Taken as a grouped averages, as in this array, ΔΔCt of gene expression changes between the species were compared and visualized using violin plot (Supplemental Figure 2B).

| **A. Heatmap ΔCt data** | | | | | | | |
| --- | --- | --- | --- | --- | --- | --- | --- |
| Species | ID | *Slc2a2* ΔCt  (t = 0) | *Slc2a2* ΔCt  (t = 10) | *Sglt1* ΔCt  (t = 0) | *Sglt1* ΔCt  (t = 10) | *Sl* ΔCt  (t = 0) | *Sl* ΔCt  (t = 10) |
| *Pteronotus parnellii* | TT18 | 6.867455213 | NA | 3.559717607 | NA | 5.844606344 | NA |
|  | TT19 | 10.2716057 | NA | 3.603201406 | NA | 4.35705189 | NA |
|  | TT23-07 | 7.700017933 | NA | 4.88196737 | NA | 5.262255404 | NA |
|  | TT21 | NA | 1.445631402 | NA | 1.445631402 | NA | 6.722156573 |
|  | TT23-08 | NA | 8.002572921 | NA | 6.662950493 | NA | 1.190077681 |
|  | TT27 | NA | 6.662950493 | NA | 8.002572921 | NA | 5.042138021 |
| *Anoura geoffroyi* | TT37 | 3.852861669 | NA | 8.625759279 | NA | -3.391762225 | NA |
|  | TT39 | 6.586765588 | NA | 6.718522559 | NA | -1.345402141 | NA |
|  | TT23-60 | 7.708034942 | NA | 8.447354221 | NA | -2.497986973 | NA |
|  | TT31 | NA | 8.401505792 | NA | 6.756257762 | NA | 0.327758008 |
|  | TT36 | NA | 5.6145058 | NA | 6.719471475 | NA | -2.2649403 |
|  | TT38 | NA | 8.7758343 | NA | 9.3002665 | NA | -1.4028583 |
|  | TT23-59 | NA | 5.584466769 | NA | 9.926680474 | NA | -1.995887726 |
| *Artibeus jamaicensis* | TT23-26 | 12.29180807 | NA | 15.63153132 | NA | -1.408687055 | NA |
|  | TT23-28 | NA | 11.75367605 | NA | 14.1567788 | NA | 0.976657957 |
|  | TT69 | NA | 11.26822646 | NA | 14.11465976 | NA | -1.624106145 |
| *Glossophaga soricina* | TT23-20 | -2.83783505 | NA | 7.975948341 | NA | -2.540237737 | NA |
|  | TT50 | NA | -3.035250005 | NA | 7.6965458 | NA | -3.4618445 |
|  | TT52 | NA | -3.071867371 | NA | 6.924639434 | NA | -3.563324866 |
|  | TT23-22 | NA | -1.994958137 | NA | 7.9596565 | NA | -1.513455477 |
| *Carollia perspicillata* | TT57 | 10.54529556 | NA | 5.551896537 | NA | -2.08255449 | NA |
|  | TT60 | 11.47327421 | NA | 5.603530026 | NA | -2.95612763 | NA |
|  | TT23-30 | 10.5643749 | NA | 6.877069929 | NA | -1.124668196 | NA |
|  | TT59 | NA | 11.39375879 | NA | 6.174707817 | NA | -2.112814101 |
|  | TT23-03 | NA | 11.40683157 | NA | 7.148294082 | NA | -1.869435076 |

| **B. Violin Plot mean ΔΔCt data** | | | | |
| --- | --- | --- | --- | --- |
| Species | ID | *Slc2a2* ΔΔCt  (t = 10) | *Sglt1* ΔΔCt  (t = 10) | *Sl* ΔΔCt  (t = 10) |
| *Pteronotus parnellii* | TT21 | -6.834061545 | 2.349068866 | 1.567518694 |
|  | TT23-08 | -1.616742454 | -2.559467781 | -3.964560198 |
|  | TT27 | -0.277120026 | 0.436678094 | -0.112499858 |
| *Anoura geoffroyi* | TT31 | 2.352285059 | -1.174287591 | 2.739475121 |
|  | TT36 | -0.434714933 | -1.211073878 | 0.146776813 |
|  | TT38 | 2.726613567 | 1.369721147 | 1.008858813 |
|  | TT23-59 | -0.464753964 | 1.996135121 | 0.415829387 |
| *Artibeus jamaicensis* | TT23-28 | -1.02358161 | -1.516871554 | -0.21541909 |
|  | TT69 | -0.538132025 | -1.474752518 | 2.385345012 |
| *Glossophaga soricina* | TT50 | -0.197414956 | -0.279402541 | -0.921606763 |
|  | TT52 | -0.234032321 | -1.051308908 | -1.023087129 |
|  | TT23-22 | 0.842876912 | -0.016291841 | 1.02678226 |
| *Carollia perspicillata* | TT59 | 0.532777238 | 0.163875653 | -0.058363996 |
|  | TT23-03 | 0.545850018 | 1.137461918 | 0.18501503 |

**Supplementary Table 8:** (A) Primer-sequences for RT-qPCR were designed targeting conserved sequences between species across exon junctions and (B) results presented as an ANOVA table from statistical analysis performed on the ΔΔCt values comparing fold change for gene expression at t=10 relative to t=0.

| **A. qPCR Primers** | | | |
| --- | --- | --- | --- |
| **Gene Symbol** | **Fragment length bp** | **5’-3’ primer sequence** | **Primer Origin** |
| *GAPDH* | 137 | F: CCT TCA TTG ACC TCA ACT AC  R: ATC TCG CTC CTG GAA GAT | This study |
| *Slc2a2* | 120 | F: CCG ACA GCC TAT TCT AGT GGC ATT G  R: TGT TGA TGG CAC CAA CTC CGA TG | This study |
| *Sglt1* | 100 | F: GTT GGA TTC TTC CTG  R: CCA GCC CCA CAA AGT | This study |
| *SI* | 126 | F: CTG GAT CCA GCA ATT TCA  R: ATC TGG CCA AAC CTT TG | This study |

| **B. Statistical Analysis** | | | | | |
| --- | --- | --- | --- | --- | --- |
| ***Slc2a2*** | | | | | |
|  | **Sum of Squares** | **DF** | **Mean Square** | **F** | **P-value** |
| **Treatment** | 30.09 | 4 | 7.523 | F (4, 9) = 1.999 | P=0.1784 |
| **Residuals** | 33.87 | 9 | 3.764 |  |  |
| **Total** | 63.97 | 13 |  |  |  |

| ***Sglt1*** | | | | | |
| --- | --- | --- | --- | --- | --- |
|  | **Sum of Squares** | **DF** | **Mean Square** | **F** | **P-value** |
| **Treatment** | 5.945 | 4 | 1.486 | F (4, 9) = 0.6146 | P=0.6630 |
| **Residuals** | 21.76 | 9 | 2.418 |  |  |
| **Total** | 27.71 | 13 |  |  |  |

| ***SI*** | | | | | |
| --- | --- | --- | --- | --- | --- |
|  | **Sum of Squares** | **DF** | **Mean Square** | **F** | **P-value** |
| **Treatment** | 8.666 | 4 | 2.167 | F (4, 9) = 0.7431 | P=0.5862 |
| **Residuals** | 26.24 | 9 | 2.916 |  |  |
| **Total** | 34.91 | 13 |  |  |  |
